# High-throughput functionalization of the *Toxoplasma* kinome uncovers a novel regulator of invasion and egress

**DOI:** 10.1101/2021.09.23.461611

**Authors:** Tyler A. Smith, Gabriella S. Lopez-Perez, Emily Shortt, Sebastian Lourido

## Abstract

Protein kinases regulate fundamental aspects of cell biology in all eukaryotes, making them attractive chemotherapeutic targets in Apicomplexan parasites such as the causative agents of malaria (*Plasmodium* spp.) and toxoplasmosis (*Toxoplasma gondii*). However, the precise roles of individual parasite kinases cannot be inferred simply from sequence identity, due to rewiring of signaling pathways and the shifting repertoire of kinases across species. To systematically examine the parasite kinome, we developed a high-throughput (HiT) CRISPR-mediated tagging strategy to endogenously label all predicted cytosolic protein kinases with a synthetic sequence encoding the minimal auxin-inducible degron (mAID) linked to a fluorophore and epitope tag. The system enables the assembly of thousands of tagging vectors from synthetic sequences in a single reaction and the pooled generation of mutants to examine kinase localization and function. We examined the phenotypes associated with kinase knock-down in 1,160 arrayed clones by replica-plating in the presence or absence of auxin and found broad defects across the lytic cycle for 109 clonal isolates, assigning localizations to 39 proteins, and associating 15 kinases within 6 distinct morphological phenotypes. The relative fitness of tagged alleles was also examined by tracking the relative abundance of individual guide RNAs as parasite populations progressed through the lytic cycle, in the presence or absence of auxin. Pooled screening had a high predictive value and differentiated between delayed and acute death. Demonstrating the value of this resource, we identified a novel kinase associated with delayed death as a novel regulator of invasion and egress. We call the previously unstudied kinase Store Potentiating/Activating Regulatory Kinase (SPARK), based on its impact on intracellular Ca^2+^ stores at key moments during the lytic cycle. Despite having a similar kinase domain to the mammalian PDK1, SPARK lacks the canonical lipid-binding domain and we find no indication SPARK positively regulates other AGC kinases, suggesting a rewiring of signaling pathways to accommodate parasite adaptations. The HiT vector screening system extends the applications of genome-wide screens into complex cellular phenotypes, providing a scalable and versatile platform for the dissection of apicomplexan cell biology.

## INTRODUCTION

Apicomplexans are widespread parasites that include the causative agents of toxoplasmosis (*Toxoplasma gondii*), cryptosporidiosis (*Cryptosporidium* spp.), and malaria (*Plasmodium* spp.). Despite their global health burden, many aspects of apicomplexan cell biology remain poorly understood. Kinases can provide important insight into parasite biology as key regulators of cellular processes that are consequently attractive drug targets ^1–3^. However, *in silico* inference of kinase function is challenging since evolutionary divergence and parasite-specific adaptations drive the rewiring of signaling pathways away from their established roles in other systems. Even broadly conserved kinase families like the cyclin-dependent kinases (CDKs) have acquired dramatic lineage-specific adaptations ^4^. The apicomplexan kinome also contains phylum-specific ROPK, FIKK, and WNG families of secreted kinases that evolved to interfere with the signaling of host cells ^5–11^. While many studies have explored the function of individual kinases in isolation, systematic approaches are needed to achieve a global view of the relative contribution of individual enzymes to parasite biology.

Recent high-throughput gene knockout screens in both *T. gondii* and *Plasmodium* spp. have provided insight into the fitness costs associated with loss of each gene for parasite growth ^12–14^. The incorporation of molecular barcodes has enabled these approaches to profile thousands of genes in a single experiment. Efforts directed specifically at the *Plasmodium berghei* kinome identified requirements of parasite progression through the sexual cycle ^15^. However, detailed high-throughput functional characterization and identification of complex cellular phenotypes beyond general fitness costs remains a critical challenge. Moreover, the lack of temporal control over the implemented gene disruption methods prevents precise timing of essential gene functions. These screens also lack information on protein localization, which can provide insight into a kinase’s molecular function.

Several genetic systems have been developed for the precise regulation of *T. gondii* gene expression, enabling the functional analysis of individual genes. Transcriptional regulation systems provide inducible transcriptional control over target genes using heterologous promoters, but regulation may require several replication cycles ^16–18^. Conditional genome rearrangements are also possible using a rapamycin-dimerizable Cre recombinase (DiCre) system, to delete entire loci or modify mRNAs for post-transcriptional regulation. DiCre systems can largely retain endogenous regulation, but recombination is irreversible and, like transcriptional regulation, its effects may be delayed for several replication cycles ^19–21^. By contrast, the auxin-inducible degron (AID) system confers post-translational regulation, often achieving complete knockdown of the target protein within one hour ^22–24^. The AID system is also reversible, maintains the target gene’s native promoter, and can be easily paired with a fluorescent protein tag. Deploying the AID system at scale could provide both subcellular localization and temporal resolution for the examination of phenotypes associated with target gene loss.

High rates of non-homologous end joining (NHEJ) typically preclude efficient homologous recombination (HR) in *T. gondii*. In fungi, mutants deficient in NHEJ have been used for genome engineering, taking advantage of their improved rates of HR ^25–29^. Deletion of *T. gondii KU80*, required for NHEJ, likewise greatly increased the efficiency of modifying endogenous loci ^30,31^. The high HR efficiency of yeast has enabled large-scale gene tagging efforts to generate arrayed strain collections for protein localization by live-cell fluorescence microscopy or interaction networks by immunoprecipitation mass spectrometry ^32–34^. Recently, similar efforts have been undertaken in mammalian systems ^35,36^ providing critical insight into the function of genes beyond their general fitness cost. In parasites, arrayed protein localization surveys have been carried out in the kinetoplastid *Trypansoma brucei*; however, no such approach has been developed for an apicomplexan parasite ^37^. Stable NHEJ-deficient strains, high rates of transfection, and the adaptation of CRISPR/Cas9-based genome engineering ^38^ make *T. gondii* ideal for the development of high-throughput tagging methods.

We present a high-throughput tagging (HiT) strategy which, paired with AID conditional knockdown in *T. gondii*, is amenable to both arrayed and pooled screening. We profiled the kinome of *T. gondii* assessing subcellular localization, and defects in cell division and the lytic cycle. Our system provides spatiotemporal resolution that led to the discovery of novel regulators of various lytic cycle checkpoints, including a previously unstudied kinase critical for invasion and egress from host cells. Our system will be a powerful tool for systematically dissecting apicomplexan cell biology.

## RESULTS

### *Development of high-throughput tagging vectors for* T. gondii

While pooled knockout screening can identify general fitness phenotypes ^12^, it fails to capture protein localization and expression levels, and the precise timing of phenotype development. To that end, we developed a high-throughput tagging (HiT) strategy that uses CRISPR-directed homologous-recombination to efficiently and site-specifically integrate exogenous sequences (payloads), such as epitope tags or regulatable elements (**Fig. 1A**). This strategy is scalable thanks to rapid cloning from entirely synthetic sequences, a modular design to introduce alternative payloads, and the flexibility to construct vectors against multiple sites in a pooled format. HiT vectors encode a gRNA targeting the 3′ end of a gene as well as two 40 bp homology regions specific to the target site. Proper HR-mediated repair of the double-stranded break integrates the synthetic sequence into the genomic locus alongside a selectable marker, eliminating the gRNA target site. Integration of the gRNA sequence, along with the rest of the construct, introduces a molecular barcode through which each mutant can be identified. Because all targeting sequences are contained within a single vector, the strategy is compatible with both pooled and arrayed screening.

**Figure 1.**
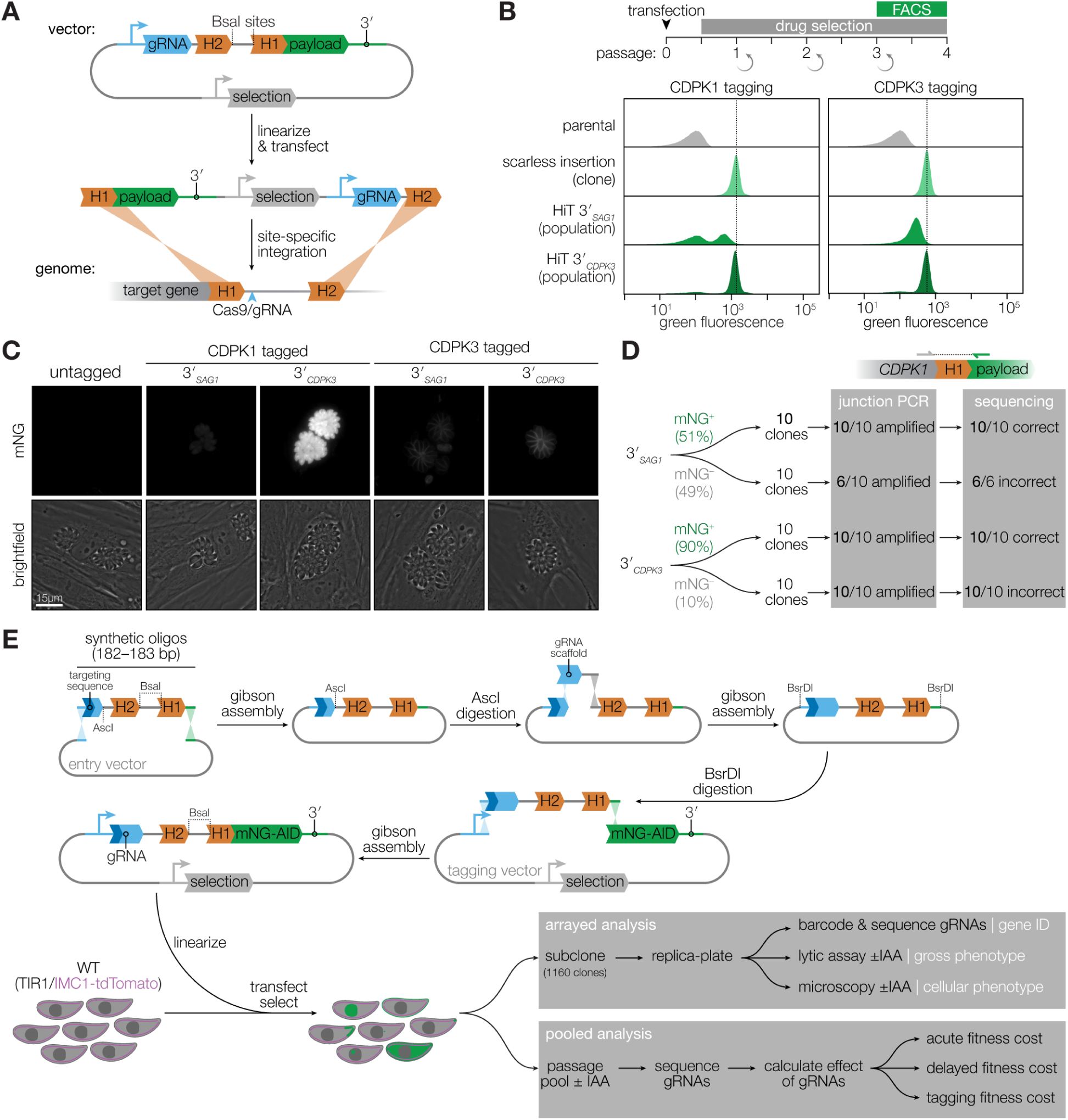
Development of high-efficiency tagging (HiT) constructs for protein-centered screening approaches. (**A**) Schematic of the high-throughput tagging (HiT) vector. The BsaI-linearized vector is cotransfected with a Cas9-expression plasmid into parasites to mediate homologous recombination of the construct into the target locus. H1 and H2 indicate 40-bp regions of homology to the target locus. Pyrimethamine selection was used to isolate successful integration events. A heterologous 3′ UTR is included in the vector following the tagging payload. (**B**) Efficiency of HiT vector tagging. Parental TIR1 parasites were transfected with HiT 3′_SAG1_ or HiT 3′_CDPK3_ vectors targeting *CDPK1* or *CDPK3* with a tagging payload encoding mNeonGreen fused to the minimal auxin-induced degron (mAID). Following selection, the populations were analyzed by flow cytometry for mNeonGreen fluorescence and compared to clonal strains carrying the same tag with no exogenous sequences (scarless insertion). Dotted line centered on the mode for the fluorescence of the scarless insertion. (**C**) Fluorescence microscopy of the tagged populations displaying the correct localization of each kinase and expression levels consistent with the flow cytometry results (B). Brightness of all mNeonGreen (mNG) images was equally adjusted. (**D**) The 5′ integration junctions of 10 mNG positive and 10 mNG negative clones from the CDPK1 HiT 3′_SAG1_ and HiT 3′_CDPK3_ populations were amplified and sequenced. Junctions were categorized according to whether they could be amplified by PCR (junction PCR) and exhibited the correct sequence at the recombination site (sequencing). (**E**) Schematic of HiT 3′_CDPK3_ vector library construction and screening strategy. Following construction and linearization of the library, the vector was co-transfected with a Cas9 expression plasmid into parasites expressing the TIR1 ligase and a fluorescent peripheral marker (TIR1/IMC1-tdTomato). Following selection, the population was analyzed by both pooled and arrayed screening.

To examine the efficiency of the HiT strategy, we designed HiT vectors to tag genes with the mNeonGreen (mNG) fluorophore fused to a minimal auxin-inducible degron (mAID). In *T. gondii* strains containing the heterologous F-box protein TIR1, addition of the auxin indole-3-acetic acid (IAA) leads to ubiquitination and proteasomal degradation of the mAID-tagged protein ^22,23,39,40^. This payload achieves both localization and regulated degradation of the gene product. As test cases, we targeted the kinases CDPK1 and CDPK3, as examples of an essential and a dispensable gene, respectively. Both kinases tolerate C-terminal tags ^41–43^. We also constructed scarless clones in which the tag was integrated into either the *CDPK1* or the *CDPK3* locus in the absence of a marker or other exogenous sequences; these control strains provide a reference to calibrate the expression of each gene from strains constructed with our HiT vectors. TIR1 parasites were transfected with HiT vectors targeting each gene together with a Cas9-expression plasmid. After selection, a majority of parasites in each population was mNeonGreen positive as assayed by flow cytometry (**Fig. 1B**). However, HiT-tagged populations were only half as fluorescent as their corresponding scarless counterparts. We hypothesized that the decreased expression may be due to the heterologous UTR selected for the vector. Replacement of the *SAG1* 3′ UTR (3′_SAG1_) with the *CDPK3* 3′ UTR (3′_CDPK3_) recovered the expression of the HiT-tagged alleles to near wild-type levels for both CDPK1 and CDPK3 (**Fig. 1B**; bottom row). The UTR-dependent changes in expression were also apparent by live-cell microscopy (**Fig. 1C**), which additionally confirmed the expected localization of the two kinases ^41,43^. Confirming the function of the tag, treatment with IAA for 24 hours resulted in complete depletion of the protein tagged in each population (**Fig. S1**).

To further characterize the source of heterogeneity within the selected populations, we isolated 10 mNeonGreen-positive and 10 mNeonGreen-negative clones from each of the tagged CDPK1 populations, and sequenced the gene-tag junction. In all cases, the mNeonGreen-positive clones from either HiT construct had the tag successfully integrated in frame. By contrast, the junction could only be amplified from 6 of the 10 negative clones originating from the population that used the 3′_SAG1_ HiT vector; in all such cases, the vector integrated into the correct locus with a frameshift mutation that precluded tag expression. Analogously, all 10 mNeonGreen-negative clones from the population that used the 3′_CDPK3_ construct had integrations marked by frameshift mutations (**Fig. 1D**). These results indicate that nearly all of the selected parasites integrated the HiT vectors in the correct locus, with a majority of parasites in the selected population harboring the tag perfectly in frame—51% of those selected with the 3′_SAG1_ construct, and 90% of those selected with the 3′_CDPK3_ construct—rates compatible with high-throughput screening.

### Generating an array of conditional mutants

Protein kinases exhibit a broad range of localizations within *Toxoplasma* to control key life cycle events such as invasion, replication, and egress ^6^. Using the HiT vector platform we chose to profile the parasite kinome. We designed constructs against 147 protein kinases and eight control genes. We included the protein kinase A regulatory subunit (PKAr), Doc2.1, a putative calmodulin, and PI-PLC based on their known or proposed roles in the lytic cycle during entry and exit from the host cell ^44–47^. CAM1, CAM2, and CAM3 were also included based on previous literature that successfully tagged these genes with the AID system ^48^. Finally, we included the hypothetical protein TGGT1_222305 as a non-kinase dispensable control ^12^. We excluded predicted ROP kinases and kinases with predicted signal peptides, based on the prediction that they are likely inaccessible to the cytosolic TIR1, and therefore of limited potential within current screen design. Previous genome-wide screens also indicated that ROP kinases are largely dispensable in cell culture ^12^.

For each gene we designed homology regions upstream and downstream from the double-stranded break, which were paired with one of three different gRNAs per gene, resulting in 465 unique constructs. To increase the possible number of gRNAs per gene, we selected gRNAs with either NGG or NAG protospacer adjacent motifs (PAMs). Homology region 1 (H1), which leads into the HiT vector tag, was defined as the 40 bp upstream of the stop codon. Because the efficiency of homologous recombination decreases with the distance of the homology regions from the double-stranded break, we only selected gRNAs that cut within a 50 bp window downstream of the stop codon ^49–52^. Homology region 2 (H2) was designed to be 40 bp in length and starts 6 bp downstream of the cutsite. This removes the gRNA cutsite after vector integration, preventing futile cycles of cutting. Potential gRNAs were scored and ranked by on-target and off-target cutting frequency using the Rule Set 2 and cutting frequency determination algorithms ^53^ and by their distance from the gene’s stop codon. These three metrics were used to create an aggregate rank, from which the three highest-ranking gRNAs were chosen for each gene.

Since the screens were performed in parallel to the analysis of UTR function, we generated libraries using the two different 3′ UTRs. The first library was created with the 3′_SAG1_ HiT vector, and the second library was created with the 3′_CDPK3_ HiT vector. Short oligos encoding the gRNA and matched homology regions were synthesized by massively parallel oligonucleotide synthesis and cloned into an entry vector. The library in the entry vector was linearized with AscI to introduce the gRNA scaffold, which exceeded the size constraints for oligonucleotide synthesis. The complete gRNA and homology regions were then cloned as a unit into the HiT vector to generate the final libraries. For the 3′_SAG1_ library, the assembled units were PCR amplified from the entry vector; however, we observed recombination leading to chimeric vectors containing gRNAs and homology regions targeting different genes. To reduce such PCR artifacts, the 3′_CDPK3_ library was cloned using a modified entry vector from which the assembled units could be excised with a restriction enzyme (BsrDI) and directly cloned into the HiT vector (**Fig. 1E**). The final strategy helps maintain library diversity while minimizing errors produced by repeated rounds of PCR amplification, thereby generating thousands of tagging vectors in a single reaction.

We carried out both arrayed and pooled screens using the constructed libraries. While pooled screening provides an easily scalable platform to compare the fitness of mutants under various conditions, arrayed screening enables precise phenotype determination through replica plating and fluorescence microscopy. To generate the clonal array, the 3′_CDPK3_ HiT vector library was transfected into parasites expressing TIR1, for conditional degradation of the AID-tagged proteins ^22–24,48^. The parasites additionally expressed a red fluorescent inner membrane complex marker (IMC1-tdTomato) to visualize the parasite ultrastructure across the replicative cycle ^54^. Following transfection and selection of the pooled 3′_CDPK3_ HiT vector library, the population was subcloned by limiting dilution. 1160 clonal strains were arrayed and passaged in 96-well plates (**Fig. 1E**). As controls, each plate was seeded with the parental TIR1 strain and a strain in which CDPK1 was endogenously tagged with the AID payload.

Sanger sequencing is cost prohibitive for the deconvolution of large numbers of clones. We therefore employed dual-indexing PCR to sequence the gRNAs from all 1160 clonal lines in a single next-generation sequencing experiment. Using barcoded forward and reverse primers, the integrated gRNAs of each well were amplified with a specific dual-index combination, allowing us to assign gRNA reads to each well of the array. A well was designated as gRNA-containing if a single gRNA had more than 100 reads. Wells containing multiple integrations or mixed populations were defined as those in which a secondary gRNA contained more than 10% the number of reads assigned to the most abundant gRNA. We were able to assign gRNA identities to 79% (917) of all wells, with the remaining 21% failing to amplify (**Fig. 2A**). 87% (796) of the amplified wells contained a single gRNA, representing 49% (228) of gRNAs in the original library and 82% (127) of the targeted genes (**Fig. 2B**). 13% (121) of amplified wells contained two or more gRNAs. To prevent confounding effects from double-tagged strains, only singly tagged clones were used for phenotype and localization assignment.

**Figure 2.**
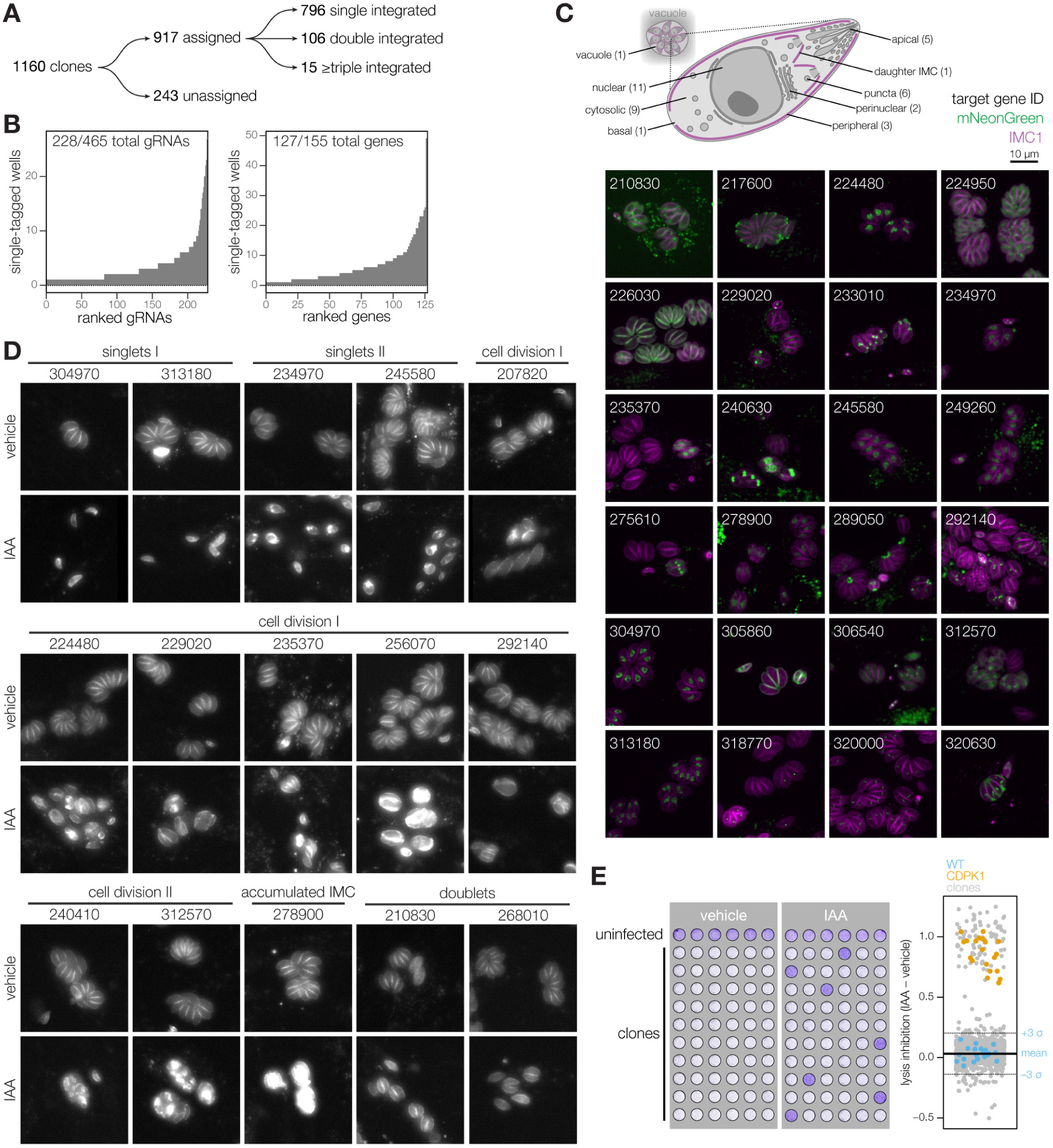
Deconvolution of protein phenotypes and localizations through high-content imaging of arrayed HiT clones. (**A**) Results from dual-indexed sequencing of the arrayed clones. A minimum of 100 reads were required to assign a given gRNA to a particular clone. Cases where a second gRNA reached >10% the abundance of the first gRNA were classified as containing multiple integrations. (**B**) Histogram showing the distribution of gRNAs and genes contained among single-integrated wells within the array. Only gRNAs and genes with an instance of at least 1 are plotted. (**C**) Distribution of localizations contained in the array; number of proteins found in each subcompartment indicated in parentheses. Localizations were assigned to a gene if half or more of single-integrated wells for that gene displayed consistent localizations. Displayed are representative confocal images of a sample of clones. mNeonGreen (green); IMC1-tdTomato (magenta). Genes are numbered based on the unique identifier from ToxoDB (e.g. TGGT1_210830, labeled 210830). (**D**) Widefield microscopy of representative clones with identified phenotypes. The IMC1-tdTomato marker is displayed for cultures treated with either vehicle or IAA for 24 h. Phenotypes were binned into 6 categories based on their similarity. (**E**) The ability of clones to lyse host cell monolayers was assayed by infecting monolayers for 72 h in the presence or absence of IAA. Intact monolayers were visualized by crystal violet staining. Normalized absorbance measurements comparing vehicle- and IAA-treated wells are graphed for each clone. Each plate contained the parental strain (WT) and an AID-tagged CDPK1 clone (CDPK1) as controls. Mean + S.D. for WT controls are shown.

### Arrayed screening to determine protein localizations and phenotypes

We employed high-content imaging to determine the localization of tagged proteins, visualize their depletion following IAA treatment, and observe the cellular consequences of their knockdown. The arrayed clones were replica-plated into 96-well glass-bottom plates containing human foreskin fibroblast (HFF) monolayers, treated with IAA or vehicle 3 hours post-infection, and imaged the following day. A clear mNeonGreen signal was observed in 29% (232) of singly tagged clones, allowing us to unambiguously determine the localization of 39 proteins, representatives of which were imaged by confocal microscopy (**Fig. 2C, Fig. S2, Table S1**). Based on the stereotypical polarized morphology of the parasites and the IMC marker, proteins could be assigned to a wide variety of subcellular compartments including nucleus (11), cytosol (9), apical end (5), cellular periphery (3), perinuclear space (2), basal end (1), daughter-cell IMC (1), parasitophorous vacuole (1), and other punctate structures (6). 36 of the 39 proteins observed showed complete depletion after 24 hours of IAA treatment. By contrast, TGGT1_322000 exhibited only partial depletion, and TGGT1_320630 and TGGT1_234950 showed no signal reduction under IAA treatment. Since TGGT1_320630 localized to the parasitophorous vacuole and TGGT1_234950 was localized to dense granules by HyperLOPIT ^55^, lack of degradation likely resulted from inaccessibility to the TIR1 machinery. Our results illustrate how the HiT screening strategy can be used to determine the localization of parasite genes in a high-throughput manner.

We were able to detect knockdown-induced phenotypes, using IMC1-tdTomato as a marker for parasite structure in the IAA-treated wells to categorize mutants based on their effects on replication and overall morphology (**Fig. 2D**). Clones bearing gRNAs against TGGT1_304970, TGGT1_313180, TGGT1_234970, or TGGT1_245580 arrested early in the lytic cycle and were mostly observed as single parasites within host cells. Degradation of the latter two additionally caused a discontinuity in the IMC marker (singlets II). Consistent with our results, a previous study categorized TGGT1_304970 as a nuclear cyclin–related kinase (TgCrk1) and demonstrated that its loss caused abnormal assembly of the daughter cell cytoskeleton and failure to progress through the cell cycle ^56^. TGGT1_234970 was also previously localized to the nucleus and categorized as the tyrosine kinase-like protein TgTKL2 ^57^. Although the function of TgTKL2 has not been determined, its similarity to the human Tousled-like kinase Tlk2 suggests it may serve a similar role promoting DNA replication ^58–60^, which would be consistent with the early arrest we observed. The two other kinases that caused an early arrest also localize to the nucleus but have not been studied in detail. TGGT1_313180 appears to be an ortholog of PRP4, which in fission yeast regulates pre-mRNA splicing and its inhibition leads to transient arrest during the gap phases of the cell cycle ^61^. By contrast, TGGT1_245580 is only conserved among coccidians (e.g. *Toxoplasma, Neospora, Hammondia, Eimeria*, and *Cyclospora*) and is predicted to be a large 2,313 residue protein with no annotated features aside from the kinase domain.

We catalogued several other phenotypes involving compromised cell division that resulted in vacuoles of smaller (cell division I) or larger (cell division II) size with aberrant IMC morphologies (**Fig. 2D**). Within this category, we identified two previously described cyclin-related kinases, TGGT1_256070 (TgCrk4) and TGGT1_229020 (TgCrk5). Consistent with our findings, knockdown of TgCrk4 resulted in major morphological abnormalities and decreased plaque size ^56^. Although the function of TgCrk5 has not been previously examined, it was previously localized to the centrocone—consistent with our observations (**Fig. 2C**)—and shown to interact with the cell cycle regulatory factor ECR1 ^62^. Strengthening the functional link between the two proteins, knockdown of TgCrk5 in our screen caused the same phenotype observed at the restrictive temperature with a temperature-sensitive allele of *ECR1* ^62^. Other cell division defects were associated with knockdown of the centrosome-associated kinases NIMA-related kinase 1 (TgNek1; TGGT1_292140), MAPK-like protein 1 (TgMAPK-L1; TGGT1_312570), and TgMAPK2 (TGGT1_207820), phenocopying the results of previously characterized temperature-sensitive or auxin-sensitive alleles ^63–66^. For TgMAPK-L1, we observe a cell cycle–dependent localization to diffuse puncta, consistent with its reported localization to the pericentrosomal matrix. For TgMAPK2, in addition to the previously reported cytosolic localization ^66^, we observe localization to paired puncta in a subset of vacuoles, which may represent cell cycle–regulated recruitment to centrosomes or early daughter cells. This localization sheds additional light into the function of TgMAPK2 and highlights an advantage of utilizing live-cell microscopy.

Cell cycle defects were also observed for several kinases that have not been previously studied. Knockdown of TGGT1_240410 resulted in a cell division II phenotype comparable to TgMAPK-L1 disruption, despite its limited conservation across closely related coccidians (*Neospora* and *Hammondia*). TGGT1_235370 exhibits a similar localization and cell division I phenotype to TgMAPK2. A cell division I phenotype was also observed following TGGT1_224480 knockdown. We show that TGGT1_224480 localizes to the nucleus, similarly to related Cdc-like kinases in mammalian cells that regulate the spliceosome ^67^, suggesting a possibly conserved function. In some cases, parasite replication arrested after a single cell cycle. Such doublet phenotypes were observed for knockdown of TGGT1_210830 or TGGT1_268010. Although conservation of TGGT1_268010 appears limited to coccidians, TGGT1_210830 is a putative ortholog of RIO kinase 1, which has been implicated in ribosomal small subunit maturation and cell cycle progression ^68^. A distinct phenotype was observed for knockdown of TGGT1_278900, which caused accumulation of the IMC marker with no specific defect in its organization. TGGT1_278900 bears homology to the yeast protein BUD32 and its *Plasmodium falciparum* orthologue complexes with other components of the EKC/KEOPS complex, suggesting broad conservation of the complex across eukaryotes ^69^.

We also screened for aggregate defects in the lytic cycle by examining monolayer clearance after knockdown. Clones were replica-plated onto fibroblast monolayers and treated for 72 hours with vehicle or IAA before quantifying monolayer integrity by measuring the absorbance of crystal violet staining (**Fig. 2E**). Changes in monolayer clearance were compared to the controls included in every plate: wild-type TIR1 parasites and an AID-tagged CDPK1 mutant that should fail to lyse the monolayer under IAA. Of all 1200 wells, 11% (129) of the wells scored three standard deviations above the average lysis inhibition score of the 20 WT negative controls, including all 20 CDPK1 positive controls. Based on this metric, these 129 clones represented 21 genes that, when conditionally knocked down, had severe defects clearing the fibroblast monolayers, including every mutant for which phenotypes had been found by high-content imaging except for TgMAPK2, which had displayed an incompletely penetrant phenotype during the time point analyzed. Importantly, clones in which ERK7 or Doc2.1 were tagged showed strong defects in monolayer clearance but displayed no defects by microscopy. This is consistent with their reported roles in invasion and egress but not replication ^46,70,71^. TGGT1_306540 shared a similar phenotypic profile to ERK7 and Doc2.1 and was previously characterized as an ethanolamine kinase (TgEK) that localized to the parasite cytosol ^72^. We similarly observed the cytosolic localization of TgEK and further established its requirement for the completion of the parasite lytic cycle. Taken together, our results highlight the power of arrayed HiT screening to capture both localization and detailed cellular phenotypes that generate new hypotheses about the function of specific kinases.

### Pooled screening distinguishes between acute and delayed-death phenotypes

Arrayed screening successfully identified phenotypes and localizations in clonal populations; however, pooled screening offers greater scalability and a more sensitive comparison of mutant fitness. To perform pooled screens, 3′_SAG1_ or 3′_CDPK3_ HiT vector libraries were transfected into TIR1 or TIR1/IMC1-tdTomato parasites, respectively. After three passages under pyrimethamine selection, the populations were split and maintained in media supplemented with either vehicle or IAA. At each passage, the parasite populations were sampled to quantify gRNA abundances by next-generation sequencing (**Fig. 3A**). The abundance of gRNAs in the pre-treatment pool correlated strongly with the abundance of gRNAs captured in the array (**Fig. S3**). We considered three possible influences on the relative abundance of a given gRNA: (i) the efficiency of vector integration, (ii) the tolerance of the locus to tagging, and (iii) the consequence of protein degradation by IAA. As expected from studies on Cas9 targeting efficiency ^53^, gRNAs designed to target sequences with NAG PAMs were less abundant than those targeting NGG PAMs in all populations post-selection (**Fig. 3B**). To regress the effect of the PAM and preserve abundance information related to the effects of tagging on the locus, we normalized the abundance for NAG gRNAs at each time point by the difference in mean abundance between NAG and NGG guides for genes previously considered dispensable ^12^ within each sample. Further, we calculated the fold change relative to the second passage, when relative gRNA abundances best reflected the mutant composition in subsequent populations. These transformations allowed us to analyze the results from both screens together, aggregating information from the relative abundance of three gRNAs across both screens for most genes.

**Figure 3.**
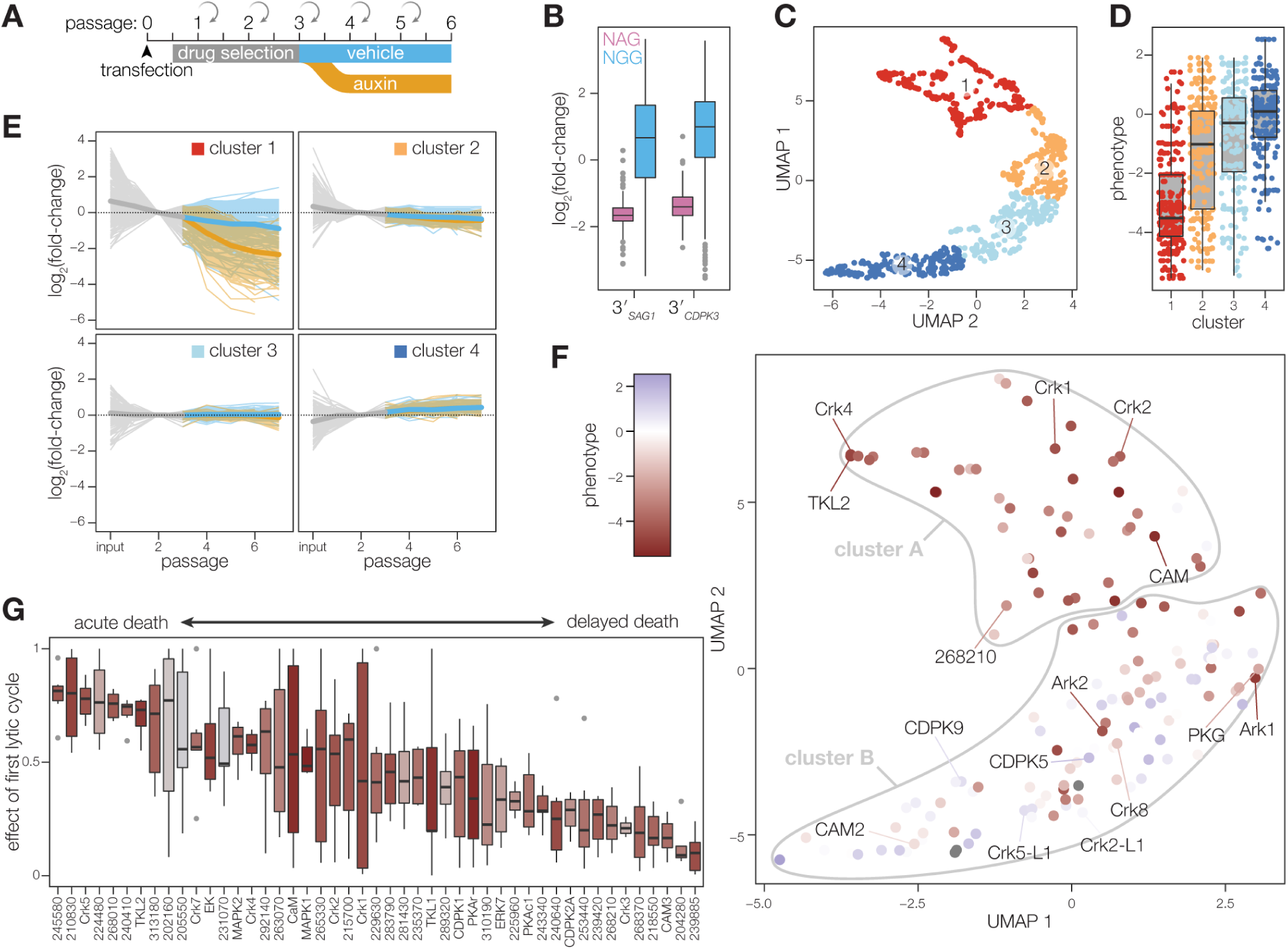
Pooled screening distinguishes between acute and delayed-death phenotypes. (**A**) Schematic of pooled screening workflow. Transfected populations were selected with pyrimethamine for three passages, after which they were split and cultured in either vehicle-or IAA-containing media. Following each lysis, parasites were split to collect samples for next-generation sequencing and continue propagating in fresh host monolayers. (**B**) Fold-change in gRNA abundance between the vehicle-treated passage 6 and the initial library, plotted by PAM type. Boxplot displays the distribution of each sample by quartiles; outliers highlighted in gray. (**C**) Relative abundances for each guide were corrected for the effect of the PAM used and fold-changes were normalized to passage 2. UMAP was used to compare guides in each screen based on their pattern of fold changes for vehicle- and IAA-treated samples. Clusters were calculated by k-means. (**D**) Comparison of phenotype scores from prior gene-disruption screen for each of the clusters in C. Boxplot displays the distribution of phenotypes in each cluster by quartiles. (**E**) Pattern of fold changes for each guide plotted by cluster. Lines colored by treatment, as in A. Bold lines are the mean for all guides in a given cluster. (**F**) Gene centroids in UMAP space based on the effects of targeting guides against each gene. Individual genes colored based on the phenotype scores from the prior gene-disruption screen. Genes were assigned to fitness-conferring (cluster A) or dispensable (cluster B) categories based on k-means clustering. (**G**) Fraction of the maximum fold-change that is explained by the first lytic cycle following IAA addition for all fitness-conferring genes (F). Genes ordered based on the magnitude of effect from acute death to delayed death.

We compared the different trajectories for gRNAs across time and IAA treatment. Using UMAP for dimensionality reduction followed by k-means clustering ^73,74^, the groupings of gRNAs generated largely agree with previously determined phenotype scores of the target genes (**Fig. 3C–D**) ^12^. Notably, cluster 1 was enriched in gRNAs that differentially dropped out of the population following IAA treatment (**Fig. 3E**). We therefore expect this cluster contains gRNAs targeting genes whose products are both fitness-conferring and successfully depleted upon IAA treatment.

To integrate the results from multiple gRNAs for the same gene, we calculated the centroid in UMAP space for gRNAs from both screens. Performing unsupervised clustering on the centroids to divide the group into two categories, we observed a cluster of 47 genes (cluster A) that was mainly composed of genes that contribute to parasite fitness (**Fig. 3F**). Considering that the arrayed screen was performed to capture acute phenotypes, it was encouraging that 18 out of the 22 genes observed to have deficits by microscopy or lytic assay were found in cluster A. 13 of the cluster A genes were simply not represented in the array, and the remaining 16 genes that were represented likely display phenotypes that are difficult to appreciate in isolation or during the brief period of kinase depletion examined in the array. One final caveat that may explain discrepancies between the two screens is the possibility that clones characterized in the array resulted from errors in homologous recombination that removed the tag—this may be a common occurrence for genes rendered hypomorphic by the presence of the tag.

Out of the four genes that appeared to have been missed by the pooled screen, for TGGT1_218720 and TGGT1_250680 a single clone was found to have a modest defect in the lytic assay. The two genes likely represent false positives in the arrayed screens, which is supported by their lack of fitness defects in our previous knockout screens ^12^. By contrast, the two other genes missed by the pooled screens (TGGT1_278900 and TGGT1_240910) are expected to be fitness-conferring ^12^. In the array, loss of TGGT1_278900 was associated with the accumulation of the IMC marker (**Fig. 2D**). Three clones of TGGT1_240910 (Doc2.1) displayed the expected lytic assay phenotype upon knockdown. Doc2.1 was originally included in the library as a control, based on prior studies ^46^. For both genes, the discrepancy between screens appears to originate from divergent results between gRNAs. Upon further analysis, each gene had a single gRNA sequence that exhibited reproducible loss in abundance when the population was treated with IAA in each screen; moreover, these were the gRNAs with optimal designs for each gene based on proximity to the coding sequence or use of an NGG PAM.

We next investigated whether the genes predicted to impact fitness in our pooled screens (cluster A) could be further categorized by the timing of gRNA loss. We sorted genes based on the fraction of the maximum effect from IAA treatment that was observed after a single lytic cycle (**Fig. 3G**). 14 of the 15 genes associated with defects by microscopy (excluding TGGT1_278900, which was missed by the pooled screens) dropped out substantially during the first passage in IAA. Among the genes that dropped out acutely but were not represented in the array, we found TGGT1_270330 (TgCrk7) and TGGT1_215700. TgCrk7 was previously reported to be essential ^56^. TGGT1_215700 has not been previously studied but is broadly conserved across eukaryotes and is homologous to phosphatidylinositol 3- and 4-kinases, which are critical for growth and cell proliferation in other organisms ^75,76^. Our results suggest that this analysis can indeed identify genes whose disruption leads to immediate and catastrophic defects in the parasite.

In contrast to the acute defects associated with some genes, gRNAs against other genes were mostly retained during the first lytic cycle or progressively lost (**Fig. 3G**). Such delayed-death phenotypes could be assigned to five genes that have been previously linked to invasion or egress: CDPK1, ERK7, PKAc1, TKL1, and CAM3 ^24,41,45,47,57,70,71^. Consistent with their delayed-death phenotypes, these genes lacked defects during the brief window selected for microscopy of the arrayed clones. Another delayed-death gene, CDPK2a, has not been previously characterized but, like CDPK1, belongs to a kinase family that has been linked to invasion and egress ^41–43,77–80^. TgCrk3 also displayed no defects by microscopy or lytic assay, consistent with the observation that knockdown resulted in reduced plaque size but normal morphology ^56^. These observations highlight that pooled screens more easily capture subtle defects that accrue over several lytic cycles and may involve the key developmental transitions that accompany egress and invasion.

### Analysis of delayed-death genes reveals two kinases that regulate invasion

We characterized four delayed-death candidates to further understand their roles in the lytic cycle (**Fig. 4A**). All four kinases were also identified as fitness-conferring in our previous genome-wide knockout screen ^12^ and remain largely uncharacterized. The four kinases exhibit different phyletic patterns based on OrthoMCL ^81^: TGGT1_204280 is conserved among several single-celled parasitic phyla, including kinetoplastids (e.g. *Leishmania* and *Trypanosoma* spp.), and amoeba (e.g. *Naegleria* spp.); TGGT1_268210 and TGGT1_239420 are conserved exclusively within the Apicomplexa; and TGGT1_239885 is restricted to coccidians, the clade of apicomplexans to which *Toxoplasma* belongs. The molecular function of all these kinases remains unclear. However, TGGT1_239420 was previously linked to the development of artemisinin resistance *in vitro* ^82^ and the *Plasmodium falciparum* ortholog of TGGT1_268210 was recently linked to regulation of protein kinase A (PKA) ^83^.

**Figure 4.**
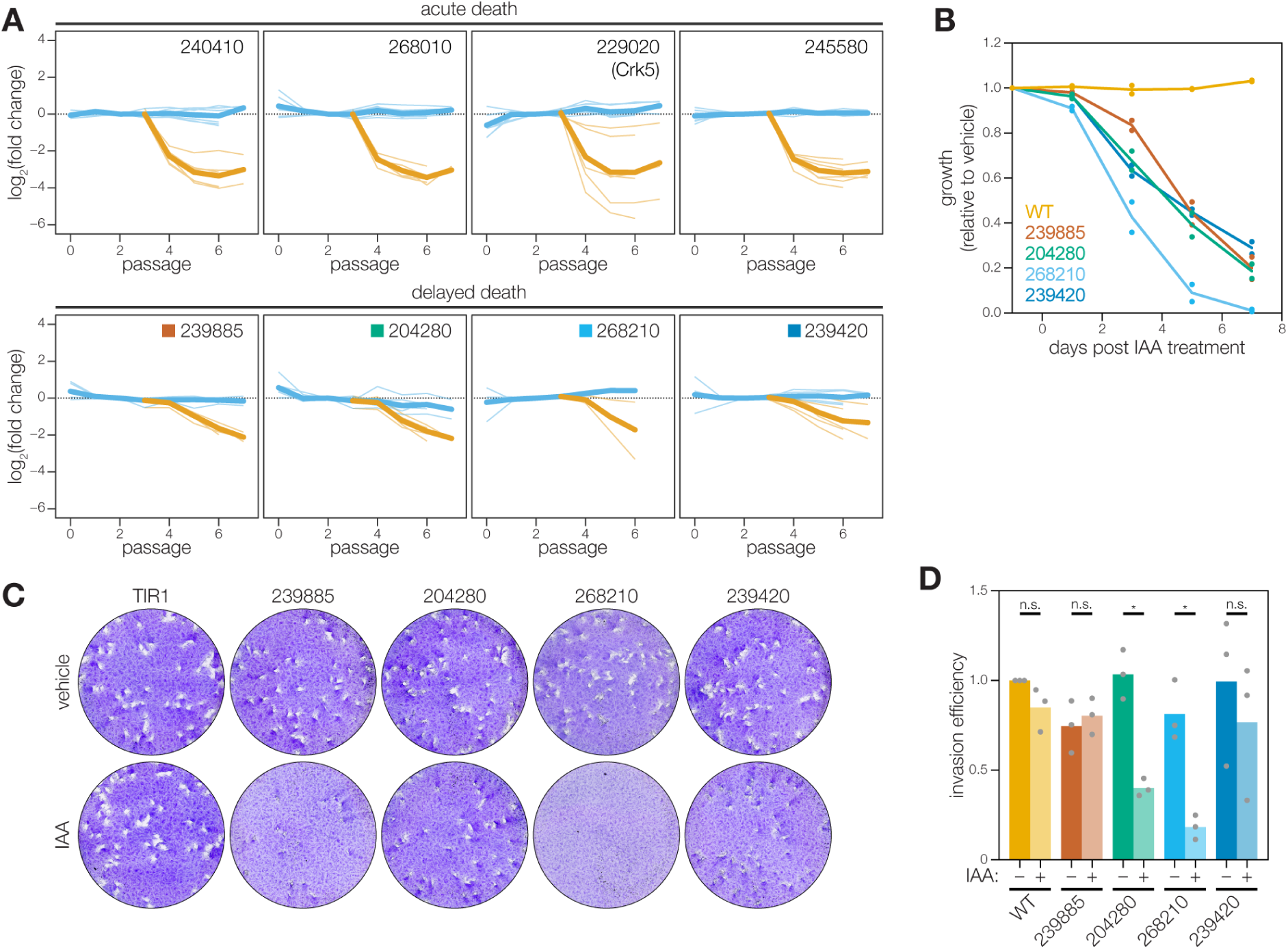
Analysis of delayed-death genes identifies two kinases that impact invasion. (**A**) Pooled screening traces of selected acute- and delayed-death genes. The fold change of individual guides in the vehicle- (blue) and IAA- (orange) treated samples are displayed. Bold lines are the mean for all guides in a given condition. (**B**) Competition assays of delayed death candidates, compared the relative growth of AID-tagged strains against a wild-type strain in either vehicle- or IAA-containing media. Following each lysis, the proportion of fluorescent parasites within each competing population was measured by flow cytometry and normalized to the vehicle control. (**C**) Plaque assays of delayed-death candidates. The ability of candidates to complete the lytic cycle was assayed by plaquing AID-tagged strains for 9 days in either vehicle- or IAA-containing media. Intact monolayers were visualized by crystal violet staining of the monolayers. (**D**) Invasion assays of delayed death candidates. AID-tagged strains grown in vehicle- or IAA-containing media for 24 h were incubated on host cells for 10 minutes prior to differential staining of intracellular and extracellular parasites. Parasite numbers were normalized to host cell nuclei for each field. Means graphed for *n* = 3 biological replicates; \**p* < 0.005, n.s. p > 0.05, Welch’s one-tailed *t*-test.

For each candidate, we rederived conditional mutants by tagging the endogenous loci with a specific 3′_CDPK3_ HiT vector. To confirm the pooled screening results, we placed each mutant in competition with TIR1/IMC1-tdTomato parasites under knockdown conditions. A second wild-type strain was used as a control for the assay. All four mutants were outcompeted by the TIR1/IMC1-tdTomato strain, demonstrating the significance of the kinases for parasite fitness (**Fig. 4B**). We also assayed the effect of knockdown for each mutant in isolation by plaque assay. Three of the mutants showed clear defects in plaque formation when grown in the presence of IAA; knockdown of TGGT1_239420 or TGGT1_239885 caused severe reductions in plaque size, while knockdown of TGGT1_268210 altogether blocked plaque formation (**Fig. 4C**). TGGT1_204280 knockdowns exhibited no plaquing defect, suggesting its effect on fitness is either subtle or only revealed in the context of wild-type coinfection. Analogously, parasites lacking TGGT1_239420 or TGGT1_239885 may be readily outcompeted by wild-type parasites due to their reduced fitness.

One possible cause of the delayed-death phenotypes could be a block in invasion. We therefore assayed the invasion efficiency of all four mutants. Both TGGT1_268210 and TGGT1_204280 displayed significant invasion defects upon knockdown (**Fig. 4D**). Since parasites formed normal plaques upon TGGT1_204280 knockdown, the invasion defect may only represent an overall delay instead of a complete block. By contrast, TGGT1_268210 knockdown caused the most severe invasion defect, which coupled with the phenotypes in plaque formation and competition assays suggests the gene plays the most critical role on the lytic cycle among the four candidates with delayed-death phenotypes. These experiments illustrate the value of examining phenotypes with temporal resolution, which helped us identify two novel regulators of invasion based on the pooled HiT screens.

### SPARK regulates the release of intracellular Ca^2+^ stores during egress and invasion

Based on its critical role during invasion, we examined TGGT1_268210 (here referred to as SPARK) in greater detail. SPARK belongs to the AGC kinase family and orthologs were found throughout the apicomplexan phylum. Although the closest mammalian orthologs of SPARK belong to the 3-phosphoinositide-dependent protein kinase-1 (PDK1) family (**Fig. 5A**), SPARK lacks the canonical C-terminal phosphoinositide-binding domain found in PDK1, similarly to related kinases in *Saccharomyces cerevisiae* and nonvascular plants ^84^. Instead SPARK and its apicomplexan orthologues possess a strongly conserved N-terminal MKXGFL motif that is absent from canonical PDK1s (**Fig. 5B**). This motif was also conserved in the most closely related orthologues of the free-living alveolates *Vitrella brassicaformis* and *Chromera velia. V. brassicaformis* and *C. velia* each also possess a second PDK1-like kinase, both of which lack the N-terminal MKXGFL motif. This suggests that gene duplication may have given rise to the SPARK clade prior to the split of the Apicomplexa from other Alveolata, which was followed by subsequent loss of the more closely related PDK1 kinase in the parasitic clade.

**Figure 5.**
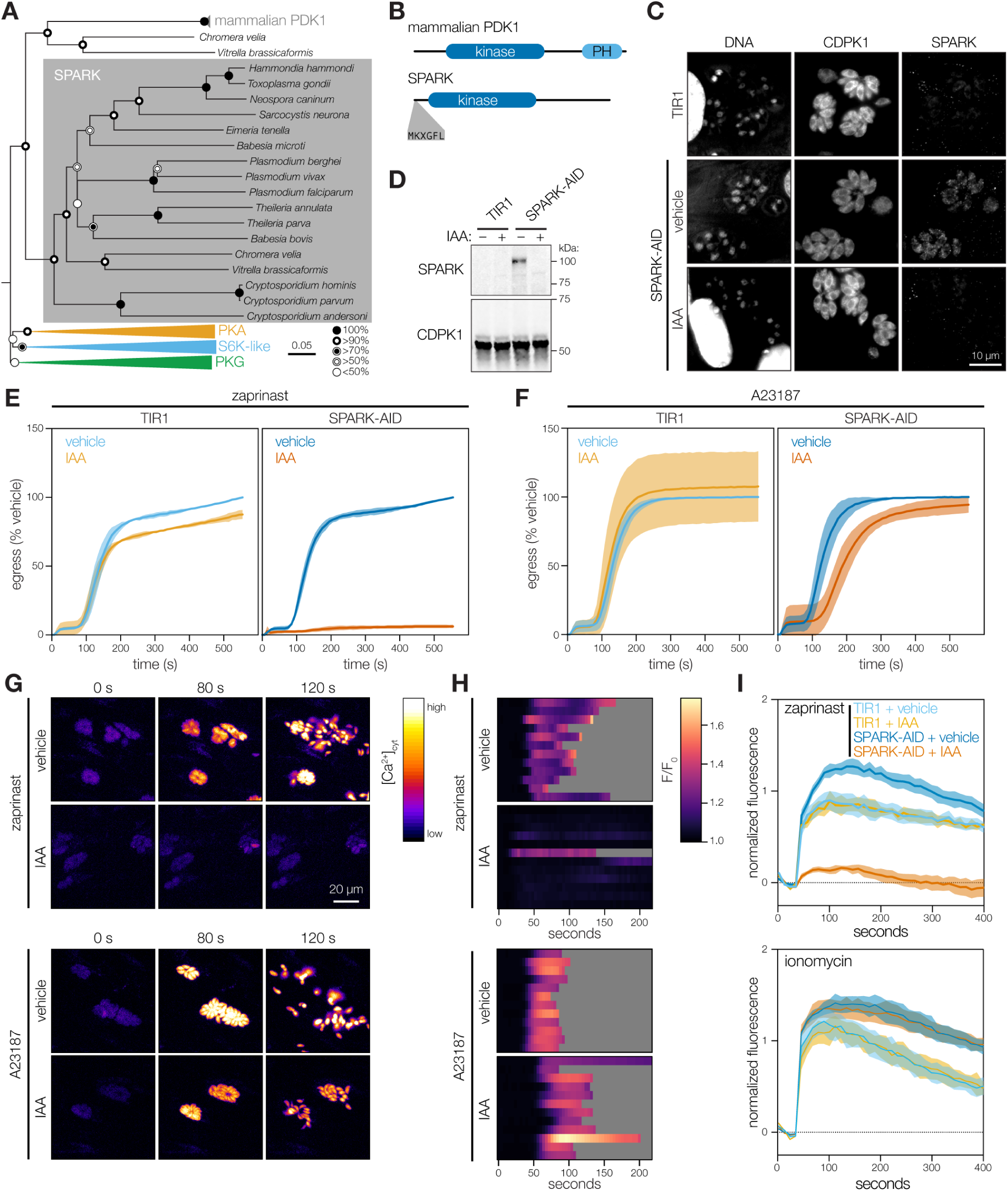
SPARK regulates egress and invasion through modulation of intracellular Ca^2+^ stores. (**A**) Neighbor-joining phylogenetic tree of kinase domains from representative apicomplexan species, along with mammalian PDK1 orthologues and related AGC kinases. Bootstraps determined from 1000 simulations. Scale indicates substitutions per site. (**B**) Models of the canonical mammalian PDK1 and the apicomplexan SPARK proteins. The kinase domains, mammalian pleckstrin homology (PH) domain, and conserved apicomplexan MKXGFL motif are shown. (**C**) SPARK-AID was visualized by immunofluorescence microscopy and immunoblotting using the V5 epitope. SPARK-AID was undetectable after 24 h of IAA treatment. Staining for CDPK1 was used to identify parasites, and nuclei were stained with DAPI. Channels adjusted equivalently across all samples. (**D**) SPARK-AID depletion, as in C, monitored by immunoblot. SPARK-AID is expected to run at 98 kDa. (**E–F**) Parasite egress stimulated with zaprinast (E) or the Ca^2+^ ionophore A23187 (F) following 24 h of treatment with vehicle or IAA. Egress was monitored by the number of host cell nuclei stained with DAPI over time. Mean ± S.D. graphed for *n* = 3 biological replicates. **(G)** Selected frames from live video microscopy of zaprinast- or A23187-stimulated SPARK-AID parasites expressing the genetically encoded Ca^2+^ sensor GCaMP6f. Parasites were grown for 24 h with vehicle or IAA prior to the stimulation of egress. **(H)** Kymographs showing normalized fluorescence per vacuole relative to the initial intensity, for 12 vacuoles per strain from the experiments in G. Gray areas represent the period following egress of the vacuole under observation. **(I)** Extracellular parasites in basal Ca^2+^ buffer stimulated with zaprinast or the Ca^2+^ ionophore ionomycin, following 24 h of treatment with vehicle or IAA. Cytosolic Ca^2+^ flux was measured in bulk as GCaMP6f fluorescence normalized to the initial and maximum fluorescence following aerolysin permeabilization in 2mM Ca^2+^.

As expected, SPARK was readily depleted following 24 h of IAA treatment in the conditional mutant, as determined by immunofluorescence microscopy and immunoblotting (**Fig. 5C–5D**, and **S4**). The protein displayed a diffuse cytosolic localization by immunofluorescence, which did not clarify its role during invasion. We carried out replication and egress assays to further characterize SPARK’s role in the lytic cycle. We did not observe a decrease in replication following 24 h of IAA treatment, narrowing the function of SPARK to the lytic cycle stages associated with parasite motility (**Fig. S5**). Since egress shares several signaling pathways with invasion, we examined the role of SPARK in this process. We induced parasites to egress from host cells by increasing cGMP levels with the phosphodiesterase inhibitor zaprinast. The resulting surge in cGMP activates PKG, which triggers release of intracellular Ca^2+^ stores ^85–90^. The Ca^2+^ ionophore A23187 was used as an alternative trigger for egress, which, despite converging with the PKG pathway at the point of increasing cytosolic Ca^2+^ concentrations, has been shown to circumvent altered guanylate cyclase activity ^91–93^. We used DAPI entry into host cells as a kinetic readout of egress ^94^. SPARK knockdown completely blocked zaprinast-induced egress (**Fig. 5E**). By contrast, SPARK appeared almost entirely dispensable for A23187-induced egress, with knockdown causing only a slight delay in egress (**Fig. 5F**). These results suggest that SPARK loss interferes with the ability of PKG to trigger intracellular Ca^2+^ release.

To confirm SPARK’s role in Ca^2+^ release, we tagged the endogenous gene via HiT vector with a mCherry-mAID epitope in parasites expressing TIR1 and the genetically-encoded fluorescent Ca^2+^ sensor GCaMP6f ^95^. Treatment of GCaMP6f/SPARK-AID parasites with IAA severely decreased levels of cytosolic Ca^2+^ fluxes in intracellular parasites following zaprinast-stimulation (**Fig. 5G & 5H**). As with egress, this phenotype could be rescued via A23187 treatment. Rises in parasite cytosolic Ca^2+^ can be caused by either release of intracellular Ca^2+^ stores or import of extracellular Ca^2+ 96,97^. To determine whether SPARK knockdown blocks intracellular Ca^2+^ release, we stimulated extracellular parasites in a buffer containing basal Ca^2+^ concentrations (∼100 nM free Ca^2+^), removing the contribution of extracellular Ca^2+^ to observed changes in GCaMP fluorescence. As observed for intracellular parasites, SPARK knockdown blocked zaprinast-induced release of intracellular Ca^2+^ stores (**Fig. 5I & S6**). By contrast, the response to the Ca^2+^ ionophore ionomycin was unchanged by SPARK knockdown, indicating that intracellular Ca^2+^ stores are intact but unresponsive to zaprinast in these parasites. These results establish SPARK as a critical regulator of the parasite intracellular Ca^2+^ store discharge that precedes both invasion and egress. Due to SPARK’s divergence from its closest mammalian ortholog PDK1, including a lack of a phosphoinositide-binding domain, we renamed the kinase Store Potentiating/Activating Regulatory Kinase (SPARK), to more aptly describe its proposed role in apicomplexan biology.

## DISCUSSION

We present a novel platform for arrayed and pooled screening in *T. gondii* that provides spatiotemporal resolution to the phenotypes of target genes. Using CRISPR/Cas9-mediated repair, we targeted 155 genes for tagging with mNeonGreen and AID. This system enabled both localization and phenotypic analysis of the kinome. By utilizing pooled and arrayed screens we identified novel regulators of various processes throughout the lytic cycle. Arrayed screening assigned subcellular localizations to 39 proteins and phenotypes to 22—a majority of which were previously unknown. Pooled screening simultaneously profiled every target gene and differentiated between acute and delayed-death phenotypes. Among the genes associated with delayed death, we identified two previously uncharacterized regulators of invasion, one of which we characterize as a novel regulator of intracellular Ca^2+^ store release we named SPARK. We demonstrate that the HiT vector platform can be deployed at scale, with minimal automation, and is amenable to any 96-well plate-based assay. HiT vector screens provide improvements on existing technologies to increase the resolution with which we can probe apicomplexan biology.

We demonstrate that HiT screens can exploit the benefits of both arrayed and pooled formats. Arrayed screens provide information for individual clones, enabling the characterization of subcellular localizations and cellular defects, even in cases when they are polymorphic or incompletely penetrant. Pooled screening, by contrast, can compare all targeted genes simultaneously and assess more subtle fitness defects that are only apparent in competitive settings. Arrayed screening for protein localization in the kinetoplastid parasite *Trypanosoma brucei* successfully assigned localizations to most protein-coding genes via high-throughput transfections and microscopy of the resulting heterogeneous populations ^37^. However, these approaches were not designed to generate clonal mutants nor to assess their phenotypes through knockdown. Pooled screens in *T. brucei, Plasmodium* spp., and *T. gondii* have examined genetic contributions to fitness in a variety of settings, but lack temporal and subcellular resolution ^12–14,98,99^. The arrayed and pooled screening approaches of the HiT vector system work in tandem to achieve protein localization and phenotypic resolution inaccessible to existing screening platforms. Further, every well of the HiT vector array is a clonally derived strain and enables unambiguous localization of the tagged species and uniform knockdown for robust phenotypic evaluation. The arrays also enable recovery of individual clones for follow-up studies of individual genes.

While false positives in the HiT screens were minimal, 29 genes previously reported to be fitness-conferring ^12^ did not score as such in our pooled HiT screens. These false-negatives likely originate from poor-quality gRNAs, inaccessibility of the TIR1 system to the tagged protein, and tagging-induced hypomorphism. The reduced abundance of low-quality gRNAs in the transfected population also caused skewed sampling of gRNAs in the array. Future screens should utilize more gRNAs per gene, avoid gRNAs targeting NAG PAMs, and isolate clones as early as possible for the arrays. Another subset of genes were either refractory to tagging or conditional degradation. Two kinases within the array could be localized but failed to respond to IAA treatment, possibly due to the inaccessibility of TIR1. We had foreseen this limitation and excluded predicted rhoptry kinases from our library due to their compartmentalization within secretory organelles. Since the HiT payload could constitute regulatory elements instead of a protein tag, transcriptional regulation could be used to target IAA-insensitive targets. Finally, several gRNAs dropped out of the vehicle-treated population even after correcting for differences between PAMs, indicating tag-induced hypomorphism. The AID system may lead to basal degradation, even in the absence of IAA, causing loss-of-function defects on target proteins and reducing the signal for microscopy ^100^. Recently developed degron systems have minimized basal degradation and may help expand the number of proteins that are observable and tolerate the degron tag ^39,101^. The modularity of the HiT vector system enables it to be adapted to a wide variety of biological questions.

We identified several previously unstudied regulators of the parasite lytic cycle. We classified microscopy-based phenotypes into six distinct categories representing defects across the replicative cycle. Knockdown of any of four nuclear kinases entirely blocked progression through the replicative cycle, causing an arrest at the single-parasite stage. Such kinases are likely regulators of critical checkpoints, such as the ortholog of TLK2, which promotes DNA replication in human cells ^58–60^. By contrast, depletion of two other kinases resulted in completion of a single replicative cycle prior to an arrest as doublets. Neither gene has been studied in *T. gondii*; however, TGGT1_210830 is a putative ortholog of RIO kinase 1, a factor critical for ribosomal maturation and cell cycle progression ^68^. This raises the possibility that, following TGGT1_210830 knockdown, defects in biogenesis cause ribosomes to become limiting after a round of parasite replication. Doublet phenotypes may therefore result from restriction of resources, rather than failed checkpoint as predicted for the singlet phenotypes. Other phenotypes observed by microscopy included abnormalities in parasite morphology. Multiple genes displayed defects similar to those previously observed for TgMAPK-L1 and TgMAPK2 ^63–66^. Aberrant morphologies allude to failure in processes such as proper daughter cell assembly or cytokinesis, rather than the complete arrest in division observed as singlet and doublet phenotypes. The visualization of cellular consequences following gene knockdown expands upon general fitness screening and enables the generation of specific hypotheses regarding gene function. Further, our observed cellular defects represent acute and lethal consequences for the parasites following knockdown, establishing these kinases as promising therapeutic targets.

The temporal resolution of the HiT screen enabled the identification of genes displaying a delayed-death phenotype. We further characterized TGGT1_268210 (SPARK) and identified it as a regulator of parasite invasion and egress through the potentiation of intracellular Ca^2+^ stores. The proposed name for the kinase reflects the role we have defined for it in the regulation of Ca^2+^ stores during the parasite lytic cycle. Conditional knockdown of SPARK demonstrated that it is critical for egress following stimulation of the cGMP pathway. SPARK knockdown blocked zaprinast-induced egress and release of intracellular Ca^2+^ stores, phenotypes that were rescued by treatment with Ca^2+^ ionophores. This phenotypic pattern mirrors the regulation of the basal accumulation of filamentous actin during parasite gliding motility, defined as F-actin flux ^102^. F-actin flux is essential for proper invasion and egress and is dependent on the Ca^2+^- activated kinase CDPK1 but independent of microneme protein secretion. Following treatment with the phosphodiesterase inhibitor BIPPO—which is thought to act in an analogous manner to zaprinast—F-actin flux was blocked by chemical inhibition of PKG and rescued by A23187. This indicates that regulation and potentiation of Ca^2+^ stores by proteins such as SPARK is a crucial event preceding F-actin mobilization, microneme protein secretion, and subsequent egress and invasion.

SPARK exhibits a phenotypic profile distinct from PKG, for which knockdown or chemical inhibition cannot be rescued by Ca^2+^ ionophore ^23,47,85^. This is consistent with previously proposed models in which PKG regulates the activity of phosphoinositide phospholipase C (PI-PLC) leading to the production of the signaling molecules diacylglycerol (DAG) and inositol trisphosphate (IP_3_) ^42,44,92,103–105^. IP_3_ stimulates intracellular Ca^2+^ release and DAG is converted to phosphatidic acid (PA). The convergence of both the PA and Ca^2+^ signals is proposed to be necessary for efficient motility and secretion of microneme and rhoptry contents ^106^. Altering the activity of the cGMP-producing guanylate cyclase (GC) by either knockdown of the guanylate cyclase itself or its accessory proteins also leads to a block in zaprinast- and BIPPO-induced egress. However, reports differ on the effect of A23187 treatment on egress following GC knockdown, ranging from a nearly normal activation of egress to a complete failure to initiate the process ^91–93^. This inconsistency makes it difficult to definitively place SPARK in the pathway; nevertheless, we favor a model that places SPARK downstream of PKG, mediating its role releasing intracellular Ca^2+^ stores.

The observed phenotypes are also inconsistent with the canonical role of PDK1 activating other AGC kinases ^107,108^. As discussed above, failure to activate PKG would block both zaprinast- and ionophore-stimulated egress ^85^. Analogously, failure to activate PKA, a negative regulator of egress, would result in premature egress ^45,47^. Instead, our study indicates that SPARK functions as a positive regulator of invasion and egress via potentiation or activation of intracellular Ca^2+^ stores. This action could be achieved via direct stimulation or activation of Ca^2+^ channels, modulation of upstream positive regulators of Ca^2+^ release such as the GC, or inhibition of a negative regulator of Ca^2+^ release such as PKA. Further work will be needed to distinguish between these models. The characterization of SPARK provides a crucial molecular handle to study the activation of intracellular Ca^2+^ stores—an event that mediates key transitions in the apicomplexan life cycle. The divergence of SPARK from its closest mammalian orthologs, in addition to its essential role in the life cycle, also highlights it as a potential therapeutic target.

The HiT vector screening system expands upon current screening platforms and enables the identification of complex cellular phenotypes. The HiT vector platform is scalable and offers the versatility to investigate many aspects of parasite biology. While we chose to target the kinome, this technology can handle much larger gene sets (∼2000 genes) in its current format. The AID system is also reversibile ^22^, which could help distinguish temporary arrests in replication from lethal disruptions to identify promising drug targets. Alternative tagging epitopes, such as split GFP, can also extend HiT screening to additional biological questions. The incorporation of additional high-throughput technologies will further expand the scope and scale of the HiT vector platform. For example, high-content imaging can be combined with *in situ* sequencing to decode the integrated gRNA of single cells within a pooled population ^109^. This technology would enable the assignment of phenotypes from morphological information in a pooled setting. The throughput and imaging capabilities of the HiT vector platform can also be increased with automation and improved microscopy, as have been employed for recent protein localization efforts in human cells ^36^. Concurrently with our work, Jimenez-Ruiz and colleagues developed an alternative strategy for high-throughput phenotypic analysis of *T. gondii*, which implemented a rapamycin-inducible split Cas9 to precisely time gene disruption. Using a parasite strain encoding indicators of actin dynamics and apicoplast segregation, they screened a library targeting 320 genes and identified two novel, essential genes required for independent steps during host cell egress and invasion. Together with the HiT screens reported herein, these technologies offer unprecedented spatiotemporal resolution to screening in *T. gondii* and are powerful tools for dissecting the biology of these ubiquitous apicomplexan parasites.

## MATERIALS & METHODS

### Parasite and host cell culture

*T. gondii* parasites were grown in human foreskin fibroblasts (HFFs) maintained in DMEM (GIBCO) supplemented with 3% inactivated fetal calf serum (IFS) and 10 μg/mL gentamicin (Thermo Fisher Scientific), referred to as D3. Where noted, DMEM supplemented with 10% IFS and 10 μg/mL gentamicin was used, referred to as D10.

### Parasite transfection

Parasites were passed through 3 µm filters, pelleted at 1000 × *g* for 10 min, washed, resuspended in Cytomix (10 mM KPO_4_, 120 mM KCl, 150 mM CaCl_2_, 5 mM MgCl_2_, 25 mM HEPES, 2 mM EDTA, 2 mM ATP, and 5 mM glutathione), and combined with transfected DNA to a final volume of 400 μL. Electroporation used an ECM 830 Square Wave electroporator (BTX) in 4 mm cuvettes with the following settings: 1.7 kV, 2 pulses, 176 μs pulse length, and 100 ms interval.

### Strain generation

Oligos were ordered from IDT. All cloning was performed with Q5 2 × master mix (NEB) unless otherwise noted. Primers and plasmids used or generated in this study can be found in (**Table S2**).

#### Scarless CDPK1-mNG-AID

The V5-TEV-mNeonGreen-AID-Ty cassette was PCR amplified from plasmid BM216 and appended with repair homology arms using primers P108 and P109. Oligos P110 and P111 were duplexed and cloned into plasmid pSS013 to create the gRNA/ Cas9-expression plasmid. The resulting gRNA/Cas9-expression plasmid was co-transfected with the repair template into TIR1 parasites ^23,24^. Following the first lysis, mNeonGreen positive clones were isolated via fluorescence-activated cell sorting. Single clones were obtained by limiting dilution and verified by PCR amplification using primers P114 and P115 and sequencing with primers P112 and P113.

#### Scarless CDPK3-mNG-AID

The CDPK3-mNG-AID scarless strain was generated as CDPK1-mNG-AID above, using primers P116 and P117 for repair template amplification, oligos P118 and P119 for assembly of the gRNA/Cas9-expression plasmid, primers P120 and P121 for amplification of the integrated tag in clonal isolates, and primers P112 and P113 for the sequencing of tag junctions in clonal isolates.

#### TIR1/IMC1-tdTomato

The primers P96 and P97 were used to PCR-amplify the sequence pTUB1_IMC1-tdTomato_DHFR to yield a repair template with homology to the 5′ and 3′ ends of a defined, neutral genomic locus ^110^. Approximately 2 × 10^7^ extracellular TIR1 parasites were transfected with 50 µg gRNA/Cas9 plasmid targeting the neutral genomic locus and 6 µg of repair template. Single clones were isolated by fluorescence-activated cell sorting into 96-well plates containing uninfected HFFs. IMC1-tdTomato positive clones were subsequently identified by microscopy and verified by PCR amplification of the neutral genomic locus using primers P98 and P99.

#### TGGT1_268210, TGGT1_204280, TGGT1_239885, and TGGT1_239420 AID-tagged lines

HiT vector cutting unit gBlocks (IDTDNA) (P122–125) were cloned via NEBuilder HiFi assembly (NEB) into the pGL015 empty mNeonGreen HiT vector backbone. HiT vectors were linearized with BsaI and co-transfected with the pSS014 Cas9-expression plasmid into TIR1 parasites. Parasite populations were selected with pyrimethamine and single clones were isolated via limiting dilution. Clones were verified by PCR amplification and sequencing of the junction between the 3′ end of the gene (primers P100-P103) and 5′ end of the protein tag (primer P104 for TGGT1_268210 and primer P105 for TGGT1_204280, TGGT1_239420, and TGGT1_239885).

#### TIR1/GCaMP6f

The primers P106 and P107 were used to PCR-amplify the sequence pTUB1_GCaMP6f_DHFR3′UTR from plasmid Genbank MT345687 to yield a repair template with homology to the 5′ and 3′ ends of a defined, neutral genomic locus ^110^. Approximately 1 × 10^7^ extracellular parasites were transfected with 25 µg gRNA/Cas9-expression plasmid BM188 and 5 µg GCaMP6f repair template. Following two rounds of fluorescence-activated cell sorting, GFP-positive clones were isolated by limiting dilution.

#### TIR1/GCaMP6f/268210-AID

The TGGT1_268210 HiT vector cutting unit gBlock (IDTDNA) was cloned into the empty V5-TEV-mCherry-AID HiT vector backbone. The HiT vector was linearized with BsaI and co-transfected with the pSS014 Cas9-expression plasmid into TIR1/ GCaMP6f parasites. Parasite populations were selected with 25 µg/mL mycophenolic acid and 50 µg/mL xanthine. Single clones were isolated via limiting dilution. Clones were verified by sequencing of the junction between the 3′ end of the gene and 5′ end of the protein tag.

### Analysis of CDPK1- and CDPK3-tagged HiT vector populations

TIR1 parasites were co-transfected with 40–50 μg of BsaI-linearized HiT 3′_SAG1_ or HiT 3′_CDPK3_ vectors and the Cas9-expression plasmid pSS014. 24 h post-transfection parasite populations were selected with 3 μM pyrimethamine. Following pyrimethamine selection populations were analyzed by flow cytometry with a Miltenyi MACSQuant VYB. Populations were imaged by microscopy using a 60x objective and an Eclipse Ti microscope (Nikon) with an enclosure maintained at 37°C and 5% CO_2_. For IAA-induced depletion experiments, intracellular parasites were treated with either 50 µM IAA or an equivalent dilution of PBS for 24 h. Following treatment, parasites were passed through a 27-gauge needle, isolated by filtration, and analyzed by flow cytometry with a Miltenyi MACSQuant VYB.

### Design and cloning of HiT vector libraries

Three gRNA constructs were designed against each gene in the 155 gene library. The 3′ end of the genomic sequences (release 36, ToxoDB.org) were scanned for gRNAs containing either NGG or NAG PAMs and cutting sites within 50 bp downstream of the stop codon. gRNAs were ranked based on predicted on-target activity and off-target activity, as determined by the Rule Set 2 and Cutting Frequency Determination calculators ^53^, respectively, and by the distance of the cutsite from the gene’s stop codon. As the Rule Set 2 calculator does not take into account efficiencies of different PAMs, the on-target scores of NAG gRNAs were penalized as predicted by the Cutting Frequency Determination calculator. These ranks were used to create an aggregate rank for gRNA selection. Initially, only the highest ranking gRNAs were selected with cutsites within 30 bp of the stop codon and with a Rule Set 2 score above 0.2. If a gene did not have 3 gRNAs meeting these criteria, the criteria were processively relaxed to allow any gRNA within 30 bp, any gRNA within 50 bp and with a Rule Set 2 score above 0.2, and finally any gRNA within 50 bp. A ‘G’ was preprended to any gRNAs that did not start with one to ensure proper RNA polymerase III initiation. Synonymous point mutations were introduced to H1 homology regions containing BsaI or AscI restriction sites, in order to prevent restriction enzyme cutting during the cloning process. The guide library was synthesized by Agilent and each oligo includes a gRNA, 40 bp homology regions, an AscI restriction site for insertion of the gRNA scaffold, and tandem BsaI sites for linearization of the final constructs, all flanked by sequences for cloning into empty HiT vectors (**Table S2**). The HiT 3′_SAG1_ and HiT 3′_CDPK3_ libraries were cloned as follows. The HiT 3′_CDPK3_ library protocol resulted in both a slight increase of correctly assembled products and greater library diversity.

#### HiT 3′_SAG1_ library

The synthesized oligo library was PCR amplified with primers P1 and P2. PCR products were cloned into the pTS018 entry vector via Gibson assembly (VWR). The gRNA scaffold was amplified from pSL001 with primers P3 and P4. The amplified scaffold was inserted into AscI-digested entry vector library via NEBuilder HiFi assembly. Finally, the assembled cutting units (containing gRNA, scaffold, homology regions, and tandem BsaI sites) were PCR amplified using primers P1 and P2 and cloned into pTS020, the mNeonGreen-AID HiT 3′_SAG1_ vector, via NEBuilder HiFi assembly. All PCR steps were performed with iProof High-Fidelity DNA polymerase (Bio-Rad) and cloning products were electroporated into MegaXDH10B T1^R^ electrocompetent cells. DNA products were isolated from liquid cultures using either a ZymoPURE II Plasmid Maxiprep Kit (Zymo Research) or a Macherey-Nagel Nucleobond Xtra Maxi Kit.

#### HiT 3′_CDPK3_ library

In order to decrease polymerase-induced errors, during construction of the HiT 3′_CDPK3_ library we replaced PCR amplification steps with digestion with the type IIS restriction enzyme BsrDI. The synthesized oligo library was PCR amplified with primers P1 and P2, as in the HiT 3′_SAG1_ library. PCR products were cloned into the pTS031 entry vector via NEBuilder HiFi assembly. The gRNA scaffold was isolated from pTS028 via BsrDI digestion and inserted into AscI-digested entry vector library via NEBuilder HiFi assembly. Finally, the assembled “cutting units” were BsrDI-digested out of the entry vector library and cloned into pGL015, the mNeonGreen-AID HiT 3′_CDPK3_ vector, via NEBuilder HiFi assembly. All PCR steps were performed with iProof High-Fidelity DNA polymerase and cloning products were electroporated into MegaXDH10B T1^R^ electrocompetent cells. DNA products were isolated from solid agar plate cultures using a ZymoPURE II Plasmid Maxiprep Kit.

### Pooled HiT vector screening

For each screen, 500 µg of the HiT vector library was linearized with BsaI-HFv2, cleaned-up using Agencourt RNAClean XP SPRI paramagnetic beads, and co-transfected with the Cas9-expression plasmid pSS014 into 5 × 10^8^ TIR1 parasites in the HiT 3′_SAG1_ screen and 5 × 10^8^ TIR1/IMC1-tdTomato parasites in the 3′_CDPK3_ screen. Parasite transfections were divided between 10 separate cuvettes. Transfected parasites were used to infect twelve and ten 15 cm^2^ dishes with confluent HFF monolayers in the HiT 3′_SAG1_ and 3′_CDPK3_ screens, respectively. 3 µM pyrimethamine and 10 µg/mL DNaseI was added 24 h later. The parasites were allowed to egress naturally from host cells, isolated by filtration, and passaged onto eight 15 cm^2^ dishes with fresh monolayers. This process was repeated for two more passages, infecting each dish with approximately 2–3 × 10^7^ parasites. Following lysis of the third passage, the population was split into three 15 cm^2^ dishes containing fresh monolayers in D10 supplemented with 50 µM IAA and three 15 cm^2^ dishes in D10 supplemented with an equal dilution of vehicle (PBS). The IAA- and vehicle-treated populations were maintained and passaged in their respective conditions for three passages in the HiT 3′_SAG1_ screen and for four passages in the HiT 3′_CDPK3_ screen. The remaining parasites (∼10^8^) at select passages were pelleted and stored at -80°C for analysis. Parasite DNA was extracted using the DNeasy Blood and Tissue kit (QIAGEN) and integrated gRNA constructs were amplified with primer P5 and barcoding primers P6–22. The resulting libraries were sequenced using a MiSeq v2 kit (Illumina) with single-reads using custom sequencing primer P23 and custom indexing primer P24.

Sequencing reads were aligned to the gRNA library. Read counts were median normalized and gRNAs in the bottom 5th percentile of the input library were removed. To account for differences in NGG and NAG PAM efficiencies, relative abundances for each gRNA were corrected for the PAM used. The PAM efficiencies were calculated by comparing the abundances of only gRNAs in the selected population that target genes identified as dispensable in a previous genome-wide knockout screen ^12^. Fold-changes were normalized to passage 2. UMAP was used to compare gRNAs in each screen based on their pattern of fold changes for vehicle- and IAA-treated samples. Clusters were calculated by k-means. Gene centroids were calculated in UMAP space and assigned to the dispensable or fitness-conferring class using k-means clustering.

### Arrayed HiT vector screening

#### Generation and passaging of array

In parallel to the pooled HiT 3′_CDPK3_ screen, single clones were isolated via limiting dilution after 4 passages of drug selection. Parasites from passage 4 were collected and sequenced as in the pooled screening experiments, using primer P5 and P25 to amplify integrated gRNA constructs. 1160 clonal isolates were arrayed into twenty 96-well plates containing HFFs. Included in each plate was a well containing the TIR1/IMC1-tdTomato parental line and a well containing the CDPK1-AID scarless strain. Arrays were passaged every 3 days with a multichannel pipette by transferring 5% of the total lysed well volume to 96-well plates containing fresh HFF monolayers. Individual wells with incomplete lysis were scraped and passed with 10% of the total well volume.

#### Arrayed widefield microscopy

Freshly lysed arrays were replica plated with 12 µL into 96- well plates of HFFs maintained in FluoroBrite DMEM (GIBCO) supplemented with 3% IFS, 4 mM glutamine, and 10 mg/mL gentamicin. Replica plates were centrifuged at 150 × *g* and 18°C for 5 min and subsequently incubated at 37°C and 5% CO_2_. At 3 h post-infection, replica plates were supplemented with either PBS or IAA to a final concentration of 50 µM. At 24 h post-IAA or PBS addition, each well was imaged at 4 adjacent fields-of-view using a 40x objective and an Eclipse Ti microscope (Nikon) with an enclosure maintained at 37°C and 5% CO_2_.

#### Arrayed lytic assays

Freshly lysed arrays were replica plated with 10 µL into 96-well plates of HFFs maintained in D3 supplemented with either 50 µM IAA or PBS. Each replica plate contained 6 uninfected control wells. Replica plates were centrifuged at 150 × *g* and 18°C for 5 min and subsequently incubated for 72 h at 37°C and 5% CO_2_. Plates were washed 1x with PBS and fixed for 10 minutes with 100% ethanol. Intact monolayers were visualized by staining the plates for 5 min with crystal violet solution (12.5 g crystal violet, 125 mL 100% ethanol, 500 Ml 1% ammonium oxalate) followed by two PBS washes, one water wash, and overnight drying. Absorbance at 590 nm was read as a measure of host cell lysis. Absorbances were normalized to the average of a plate’s uninfected control wells.

#### Dual-indexed sequencing of arrays

Parasites were harvested from 100 µL of fully lysed wells by centrifugation at 1000 × *g* and 18°C for 10 min. Pellets were each resuspended in 25 µL of lysis buffer (1x Q5 buffer supplemented with 0.2 mg/mL Proteinase K) and lysed using the following conditions: 37°C for 1 h, 50°C for 2 h, and 95°C for 15 min. Guides were PCR-amplified from gDNA, with each well utilizing a unique combination of i7 index primers (P26–65) and i5 index primers (P66–95). PCR products from an individual plate were pooled and gel extracted using a Zymoclean Gel DNA Recovery Kit and subsequently pooled at equimolar ratios for sequencing. The final PCR product pool was sequenced with a MiSeq v2 kit (Illumina) using dual-indexed single-reads with primers P23 and P24. Sequencing reads were aligned to the gRNA library.

### Confocal microscopy of select clones

Parasites were inoculated into 96-well glass-bottom plates (Cellvis) of HFFs maintained in FluoroBrite DMEM supplemented with 10% IFS, 4 mM glutamine, and 10 mg/mL gentamicin. At 24 h post-infection wells were imaged using an RPI spinning disk confocal microscope maintained at 37°C and 5% CO_2_. The parental TIR1/IMC1-tdTomato strain was also imaged under identical conditions to control for background fluorescence.

### Plaque assays

Parasites were inoculated into 6-well plates of HFFs maintained in D10 and either 50 µM IAA or PBS and allowed to grow undisturbed for 9 days. Plates were washed with PBS and fixed for 10 min at room temperature with 100% ethanol. Staining was performed for 5 min at room temperature with crystal violet solution, followed by two washes with PBS, one wash with water, and drying overnight.

### Competition assays

Freshly lysed strains were filtered through 5 µm filters and spun-down at 1000 x *g* and 18°C for 10 min. Pellets were resuspended in D10 and counted. T12.5s of HFFs were infected with 1.5 × 10^6^ parasites of the TIR1/IMC1-tdTomato strain and 1.5 × 10^6^ parasites of the competitor strain. At 24 h post-infection the media of each flask was changed with fresh D10. Populations were assayed by flow cytometry following host cell lysis, and each population was passed to two wells of a 6-well plate. At 24 h post-infection the media of one well per strain was changed to D10 and vehicle (PBS) and the media of the second well changed to D10 and 50 µM IAA. Populations were passed and maintained in these conditions every two days for four passages. Following each lysis the populations were assayed by flow cytometry with a Miltenyi MACSQuant VYB. The fraction of the population that was tdTomato negative was represented as a ratio of the [percent tdTomato negative of the IAA sample]/[percent tdTomato negative of the vehicle sample] and was normalized to the initial fraction pre-splitting into +/- IAA media.

### Invasion assays

Freshly lysed strains were each passed to two flasks of HFFs containing D10 media. At 3 h post-infection one flask was supplemented with vehicle (PBS) and the second flask was supplemented with IAA to a final concentration of 50 µM. At 27 h post-infection each flask was syringe-lysed and filtered through 5 µm filters. Parasites were spun down at 1000 x *g* and 18°C for 10 min. Pellets were resuspended in invasion media (HEPES-buffered DMEM without phenol red) supplemented with 1% IFS to a concentration of 1 × 10^6^ parasites/mL. 200 µL of each parasite solution was added to 3 wells of a clear-bottom 96-well plate containing HFFs. The plate was spun at 290 x *g* and room temperature for 5 min. Plates were incubated for 10 min at 37°C to stimulate invasion. Following incubation wells were fixed with 4% formaldehyde and extracellular parasites were stained with mouse anti-SAG1 antibody ^111^. All parasites were stained by permeabilizing with 0.25% TritonX-100 and staining with guinea pig anti-CDPK1 (Covance, ^112^). Cells were subsequently stained with anti-guinea pig Alexa594 antibody (Invitrogen), anti-mouse Alexa488 antibody (Invitrogen), and Hoechst 33258 (Santa Cruz Biotechnology) nuclear dye. Samples were imaged using a Biotek Cytation3 imaging multimode reader. The number of invaded parasites per field of view was manually counted and normalized to the number of host cells in the same area. The final invasion efficiency for each replicate was normalized to the invaded parasites per host cell nuclei of the parental TIR1 vehicle sample.

### Phylogenetic analysis of SPARK

SPARK homologs were identified by BLAST search against representative apicomplexan genomes. Protein kinase domains were obtained from EupathDB based on their annotation with Interpro domain IPR011009. Sequences were curated for *Theileria* spp., *Cryptosporidium parvum, Cryptosporidium hominis, Sarcocystis neurona, Vitrella brassicaformis*, and *Chromera velia* to correct errors in the gene model. Domains from the nearest human, mouse, and macaque orthologues (as determined by BLAST) were used as outgroups. Individual domains were extracted from each sequence and aligned using ClustalX2, and the phylogenetic tree was generated by neighbor-joining. Visualizations were generated using FigTree (v1.4.4).

### Immunoblotting

TIR1 and SPARK-AID parasites were grown in D10 for 3 h before being treated with either 50 µM IAA or vehicle (PBS). After 24 h of IAA or vehicle treatment, parasites were scraped, passed through 27-gauge needles, and isolated via filtration. Parasite pellets were resuspended in lysis buffer (0.8% IGEPAL-CA630, 0.25 U/µL benzonase, and 2x Halt Protease Inhibitor Cocktail in PBS) and incubated on ice for 15 min. Lysates were combined with 1x Laemmli buffer (diluted from 5x buffer containing 10% SDS, 50% glycerol, 300 mM Tris HCl pH 6.8, 0.05% bromophenol blue) with 1% final volume β-mercaptoethanol and boiled for 10 min. Samples were run on a 7.5% SDS-PAGE gel (BioRad) and transferred overnight onto a nitrocellulose membrane in transfer buffer (25 mM Tris-HCl, 192 mM glycine, 20% methanol). Blocking and all subsequent antibody incubations were performed in 5% milk in TBS-T (20 mM Tris, 138 mM NaCl, 1 L PBS, 0.1% Tween-20). The blot was incubated for 1 h at room temperature followed by incubation with the mouse anti-V5 primary antibody (R960-25, Invitrogen) for 1 h rocking at room temperature. The blot was washed three times with TBS-T and incubated for 1 h with the anti-mouse secondary antibody (LI-COR). Following imaging the blot was incubated overnight at 4°C with the guinea pig anti-CDPK1 primary antibody (Covance, ^112^). The blot was washed three times with TBS-T, incubated for 1 h at room temperature with anti-guinea pig secondary antibody (LI-COR), washed 3x with TBS-T, and imaged. Imaging was performed using a LI-COR Odyssey CLx.

### Immunofluorescence assays

Parasites were inoculated onto coverslips containing HFFs and after 3 h were treated with either 50 µM IAA or PBS. At 24 h post-IAA addition, intracellular parasites were fixed with 4% formaldehyde and permeabilized with 0.25% Triton X-100 in PBS. Nuclei were stained with Hoechst 33342 or Hoescht 33258 (Santa Cruz) and coverslips were mounted in Prolong Diamond (Thermo Fisher). V5 was detected using a mouse monoclonal antibody (R960-25, Invitrogen). CDPK1 was detected using a guinea pig-derived polyclonal antibody (Covance) ^112^. Primary antibodies were stained with anti-guinea pig and anti-V5 Alexa-Fluor-labeled secondary antibodies (Invitrogen). Images were acquired with an Eclipse Ti microscope (Nikon) using the NIS elements imaging software.

### Replication assays

Parasites were inoculated onto coverslips containing HFFs and after 3 h were treated with either 50 µM IAA or PBS. At 24 h post-IAA addition, intracellular parasites were fixed, permeabilized, and stained as described under “Immunofluorescence assays”. For each sample, multiple fields of view were acquired with an Eclipse Ti microscope (Nikon) and the number of nuclei per vacuole were calculated from 100 vacuoles. Results are the mean of three independent experiments.

### Egress assays

Egress was quantified in a plate-based manner. HFF monolayers in a clear bottomed 96-well plate infected with 7.5 × 10^4^ parasites per well of parental or SPARK-AID for 3 h were treated with 50 µM IAA or PBS for an additional 24 h. Before imaging, the media was exchanged for FluoroBrite supplemented with 10% IFS. Three images were taken before zaprinast (final concentration 500 μM) or A23187 (final concentration 8 µM) and DAPI (final concentration 5 ng /mL) were added, and imaging of DAPI-stained host cell nuclei continued for 9 additional minutes before 1% Triton X-100 was added to all wells to determine the total number of host cell nuclei. Imaging was performed at 37°C and 5% CO_2_ using a Biotek Cytation 3 imaging multimode reader. Results are the mean of three wells per condition and are representative of three independent experiments.

### Live cell microscopy of GCaMP6f-expressing parasites

To capture egress, SPARK-AID parasites were grown in HFFs in glass-bottom 35 mm dishes (Ibidi) for 3 h at which point they were treated with either 50 µM IAA or PBS for an additional 24 h. Parasites were stimulated to egress with 500 μM zaprinast or 8 µM A23187 in Ringer’s buffer prepared without Ca^2+^ (155 mM NaCl, 3 mM KCl, 1mM MgCl_2_, 3 mM NaH_2_PO_4_, 10 mM HEPES, 10 mM glucose) and supplemented with 1% BSA (w/v) and recorded every 4 s for 220 s using an Eclipse Ti microscope (Nikon) with an enclosure maintained at 37 °C and 5% CO_2_.

### Extracellular ionomycin treatment of GCaMP6f strains

Parasites were passed to 15 cm^2^ dishes containing HFFs and after 6 h treated with either 50 µM IAA or vehicle (PBS). Following 24 h of IAA or vehicle treatment parasites the cells were washed once with PBS. The media was changed to cold Ringer’s Basal Ca^2+^ (155 mM NaCl, 3 mM KCl, 1 mM MgCl_2_, 3 mM NaH_2_PO_4_, 10 mM HEPES, 10mM glucose, 250 µM EGTA, 112 µM CaCl_2_) and parasites were harvested and isolated via syringe lysis and filtration. Parasites were spun-down and washed once with Ringer’s Low Ca^2+^ and resuspended to 1 × 10^7^ parasites/ mL. 100 µL of parasite suspension was added to a clear-bottom 96-well plate and incubated on ice for 5 min. Fluorescence was read with an excitation wavelength of 485 nm and an emission wavelength of 528 nm every 10 s in a BioTek Cytation 3. At 30 s 50 µL of 3x zaprinast (100 µM final concentration), ionomycin (1 µM final concentration), or vehicle (DMSO) was added and fluorescence readings were taken for an additional 6 min. 50 µL of 4x aerolysin (3 µg/mL final concentration) and CaCl_2_ (2 mM final concentration) was added and the assay plate was incubated at 37 °C for 10 min. Fluorescence was read every 1 min for 40 min. The assay plate was shaken for 1 s before each read. The non-fluorescent TIR1 strain was used to perform a baseline subtraction of background fluorescence. Baseline-subtracted values were normalized to initial fluorescence pre-stimulation and maximum fluorescence post-aerolysin treatment.

## ACKNOWLEDGEMENTS

We thank Bingbing Yuan for bioinformatics advice; L. David Sibley for the TIR1 strain; Wendy Salmon and the W.M Keck BIological Imaging Facility for confocal microscopy support; Peter Reddien for use of the Illumina MiSeq; Benjamin Waldman, Alice Herneisen, Elizabeth Boydston, Chris Giuliano, Alex Chan, Saima Sidik, and Benedikt Markus for technical support in generation of the array; EuPathDB and all contributors to this resource. This work was supported by funds from a National Institutes of Health grant (R01AI144369) to S.L. and a National Science Foundation Graduate Research Fellowship (2018259980) to T.A.S.

