## Supplementary Information for "High-throughput functionalization of the *Toxoplasma* kinome uncovers a novel regulator of invasion and egress"

### SUPPLEMENTAL TABLES

**Table S1.** Combined results from the HiT screens summarizing data from arrayed and pooled analyses.

**Table S2.** Oligos and plasmids used in this study.

### SUPPLEMENTAL FIGURES

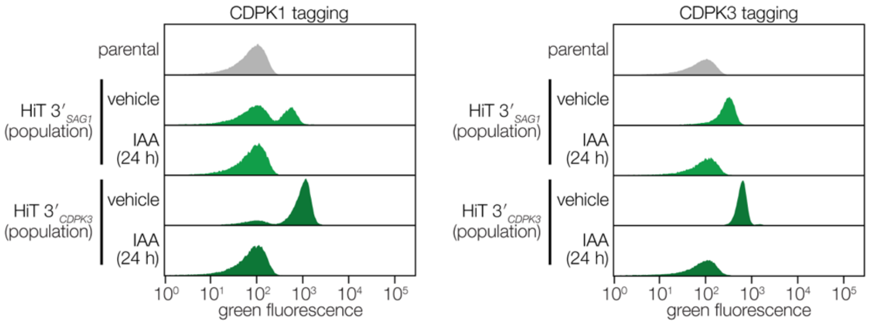


**Figure S1. HiT vector-transfected populations are IAA-responsive.** IAA regulation of HiT vector-transfected populations. Transfected populations were treated with IAA or PBS 3 hours post-infection. After 24 hours of IAA or PBS treatment populations were analyzed by flow cytometry.

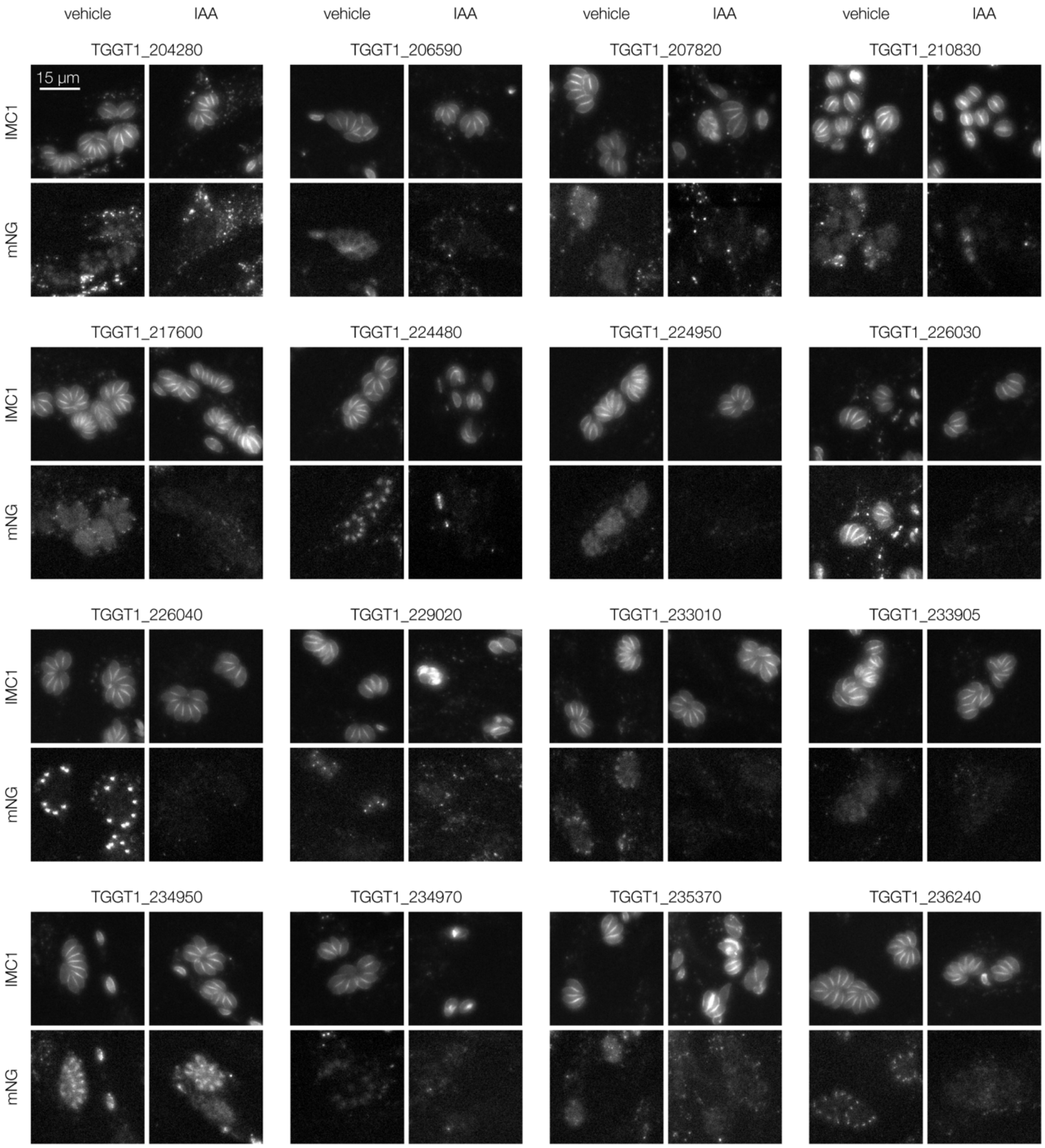
vehicle
IAA
TGGT1_204280
TGGT1_206590
TGGT1_207820
TGGT1_210830
15 µm
IMC1
mNG
TGGT1_217600
TGGT1_224480
TGGT1_224950
TGGT1_226030
TGGT1_226040
TGGT1_229020
TGGT1_233010
TGGT1_233905
TGGT1_234950
TGGT1_234970
TGGT1_235370
TGGT1_236240

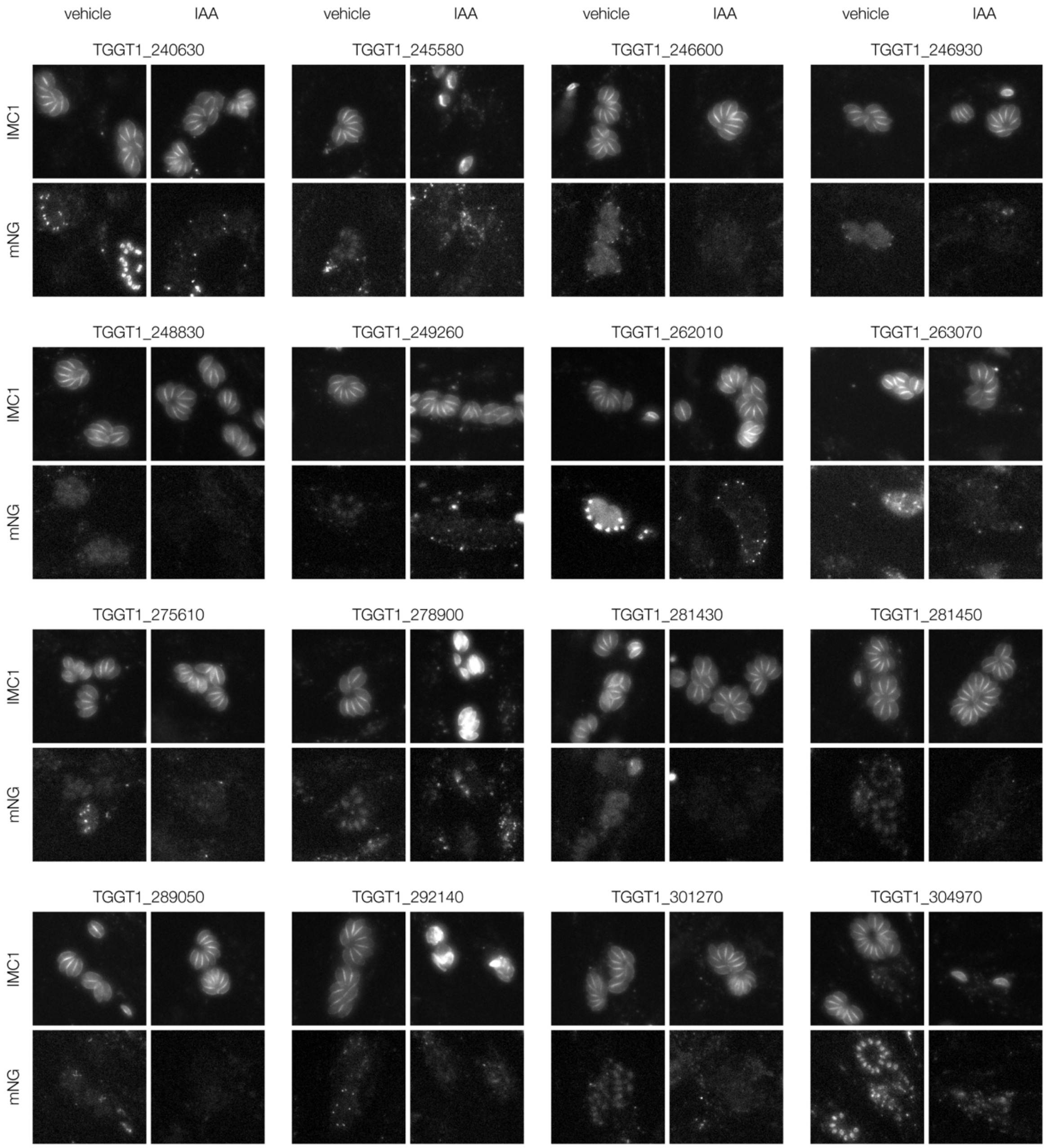
vehicle
IAA
vehicle
IAA
vehicle
IAA
vehicle
IAA
TGGT1_240630
TGGT1_245580
TGGT1_246600
TGGT1_246930
IMC1
mNG
TGGT1_248830
TGGT1_249260
TGGT1_262010
TGGT1_263070
IMC1
mNG
TGGT1_275610
TGGT1_278900
TGGT1_281430
TGGT1_281450
IMC1
mNG
TGGT1_289050
TGGT1_292140
TGGT1_301270
TGGT1_304970
IMC1
mNG

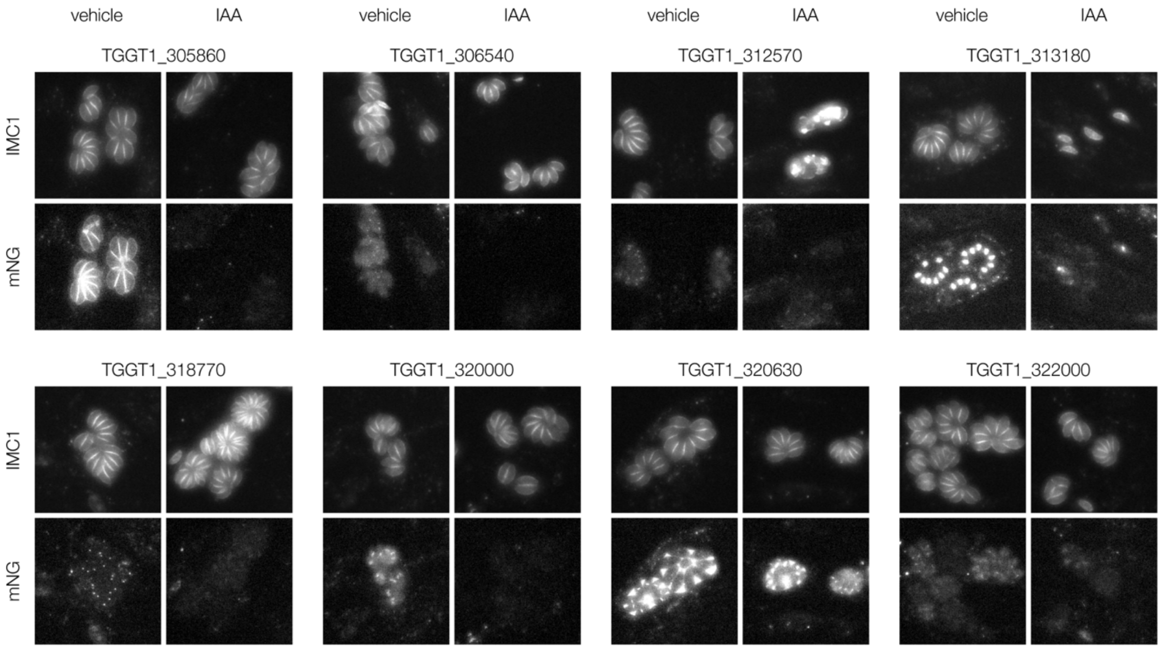


**Figure S2. Representative images of assigned localizations within the array by widefield microscopy.** Widefield microscopy of representative clones with assigned localizations. Localizations were assigned to a gene if half or more of single-integrated wells for that gene displayed consistent localizations. The maximum IMC1-tdTomato and mNeonGreen markers are displayed for cultures treated with either IAA or vehicle for 24 hours.

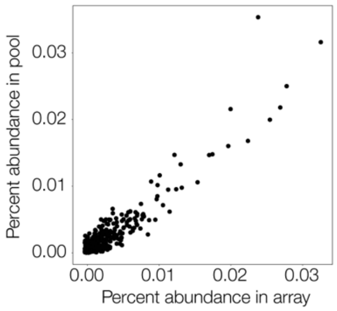


**Figure S3. Abundance of gRNAs captured in the array correlates strongly with their presence in the pooled population.** Percent abundance of each gRNA captured in the array versus the percent abundance of each gRNA in the pooled population from which they were subcloned. Spearman correlation coefficient = 0.77.

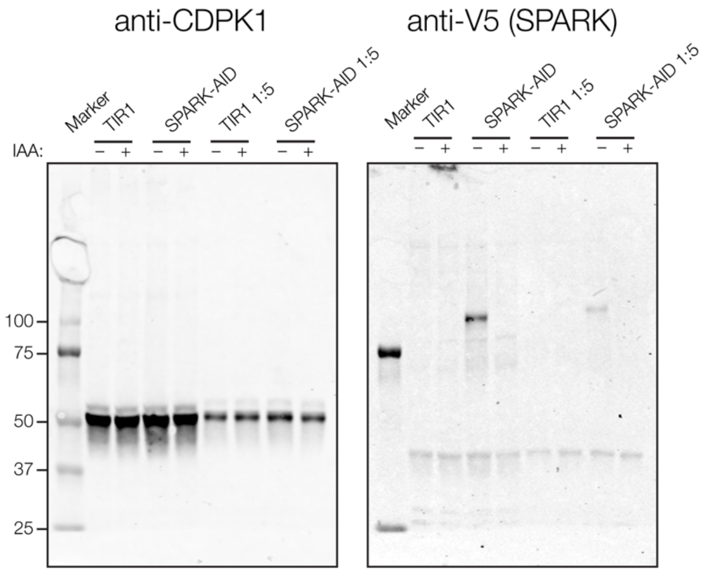


**Figure S4. SPARK-AID depletion monitored by immunoblot.** Full immunoblot from Figure 5D. SPARK-AID was detected using an anti-V5 antibody and CDPK1 was used as a loading control.

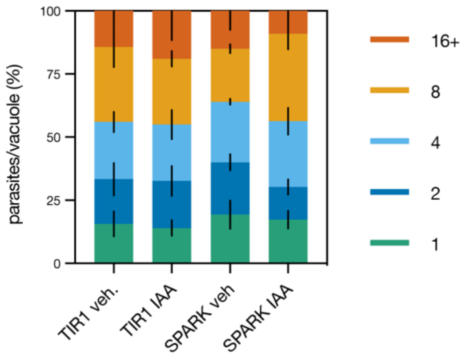


**Figure S5. Knockdown of SPARK does not lead to a replication defect.** Replication assay of SPARK-AID parasites. Parasites were treated with either IAA or vehicle at 3 hours post-invasion and incubated for 24 hours. Parasites were stained and imaged as 4x4 fields. The number of parasites per vacuole were counted until 100 vacuoles had been counted for a given sample. (N=3).

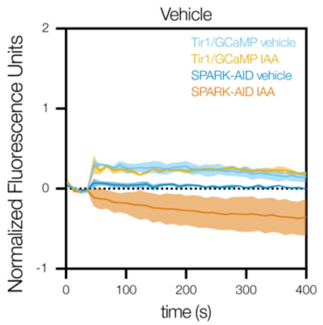


**Figure S6. Vehicle-treated extracellular GCaMP6f/SPARK-AID parasites in basal $Ca^{2+}$ buffer.** Vehicle trace from **Figure 5I**. Extracellular parasites were treated with vehicle and the fluorescence response was quantified and normalized as previously described.
